# Behavior-relevant top-down cross-modal predictions in mouse neocortex

**DOI:** 10.1101/2023.04.03.535389

**Authors:** Shuting Han, Fritjof Helmchen

## Abstract

Animals rely on predicting their environment and the consequences of their actions to adapt to a constantly changing world. The predictive coding hypothesis proposes that the brain generates predictions and continuously compares them with bottom-up sensory inputs to guide behavior. However, how the brain reconciles conflicting top-down predictions and bottom-up sensory information during behavior remains unclear. To address this question, we simultaneously imaged neuronal populations in the mouse somatosensory cortex and the posterior parietal cortex during an auditory-cued texture discrimination task. After mice learnt the task with fixed tone-texture matching, mismatched pairing caused conflicting tone-based texture predictions and actual texture inputs. When top-down interaction was dominant, texture representations in both areas were modified and mice decided based on the predicted rather than actual texture, whereas dominant bottom-up interaction corrected the representations as well as behavioral choice. Our findings provide evidence for hierarchical predictive coding in the mouse neocortex and open new avenues for understanding higher cognitive functions.

## Introduction

Predictive processing has long been an attractive theory of the mind. This theory states that the brain is organized hierarchically, with predictions being generated in high-level areas, such as frontal and posterior association areas, and passed down to lower areas in a top-down fashion. Mismatched sensory inputs that do not fit the predictions in this scheme introduce bottom-up flow that represents prediction errors^1^. Despite the computational attractiveness of this model, it remains elusive how it is implemented in the brain. While reward prediction and reward prediction errors have been studied extensively for the dopamine system and in frontal cortices^2–5^, sensory prediction in the neocortex is much less understood. Sensory prediction often originates from prior experience, typically based on learnt associations with other sensory cues. It has been observed in many sensory modalities including vision, sound, taste and touch^6–10^. Such predictions can increase the encoding speed and reduce the neural response to expected sensory inputs in primary sensory areas^6,8^, facilitating decisions and behavioral output. Strong sensory predictions can also modify perception, in extreme cases causing hallucination^11–13^.

One challenge to study sensory prediction is to simultaneously observe bottom-up and top-down information. Studies targeting long-range projection axons as a proxy for top-down inputs to local populations have demonstrated that such pathways can indeed modulate sensory perception and decision making^6,9,14^. However, studies that focus on how neuronal populations along the brain hierarchy represent and transform information, as well as how they communicate with each other, have started only recently^15–20^. These studies discovered, for example, that top-down and bottom-up information are channeled through separate activity subspaces^15,16^, and that the communication channels are shaped by experience or learning, especially the top-down subspace^18,19,21^. Despite these insights, it is still unknown how top-down predictions and bottom-up sensory inputs interact during behavior and affect behavioral outputs, particularly when they are in conflict.

A key area for routing primary sensory information during active behaviors is the posterior parietal cortex (PPC). PPC is densely interconnected with primary sensory areas such as the visual cortex, the somatosensory cortex, and the auditory cortex, as well as frontal areas such as the orbitofrontal cortex and the anterior cingulate cortex, and the associative subdivision of thalamus^22^. PPC is involved in a wide range of functions including multisensory integration, decision making, working memory, and navigation^22^. In particular, PPC has been shown to integrate tactile, visual and auditory information in rodents^23–25^, and to route relevant sensory information to frontal areas during active behaviors^26–28^. Of particular relevance to our study is that different subdivisions of PPC engage differentially in the processing of distinct stimulus modalities: we recently showed that the rostrolateral area (PPC-RL) is activated together with the primary somatosensory (S1) area during texture discrimination, whereas the anterior area (PPC-A; located medially to PPC-RL) activates in conjunction with auditory cortex areas in an auditory discrimination task^26^. These PPC areas are critical for further transforming sensory information into decisions^26,29–31^, making them attractive candidate high-level areas for studying predictive processing.

Here, we aim to better understand how cortical areas along the cortical hierarchy interact when sequential stimuli from two modalities (auditory and tactile) provide task-relevant information. In this case, repeatedly matching specific pairs of auditory-tactile stimuli allows the animal to form predictions about the second stimulus. It is then especially interesting to reveal how regional neural representations and cross-areal interactions are affected when conflicts between predictions and sensory inputs are imposed. For this study, we focused on the barrel field in S1 and PPC subdivisions as representative areas along the hierarchy. We designed a behavioral task with cross-modal sensory predictions (by training mice on matched tone-texture sequence pairs), and then introduced prediction errors by occasionally presenting mismatched tone-texture pairs. We used the behavioral choices of mice as a proxy for their perceptual representations. We found that during mismatch, predictions could modify sensory encoding in both S1 and PPC, and that such representational changes were also reflected in the behavioral choice. Moreover, these changes only occurred when the top-down prediction from the relevant subdivision of PPC-A was dominant, whereas strong bottom-up information from S1 could correct both sensory encoding and behavioral outcome. These results demonstrate the impact of predictions on sensory encoding and suggest that the dynamic interaction between top-down and bottom-up information shapes sensory information and affects perceptual choice.

## Results

### An auditory-cued texture discrimination task

To study sensory prediction, we developed an auditory-cued texture discrimination task for mice. Mice were trained to discriminate two textures (rough vs. smooth), and each texture was associated with a distinct auditory tone (10 kHz vs. 18 kHz) before texture onset. Each tone-texture sequence led to a reward from one of the two lick ports (Fig. 1a-b).

**Figure 1.**
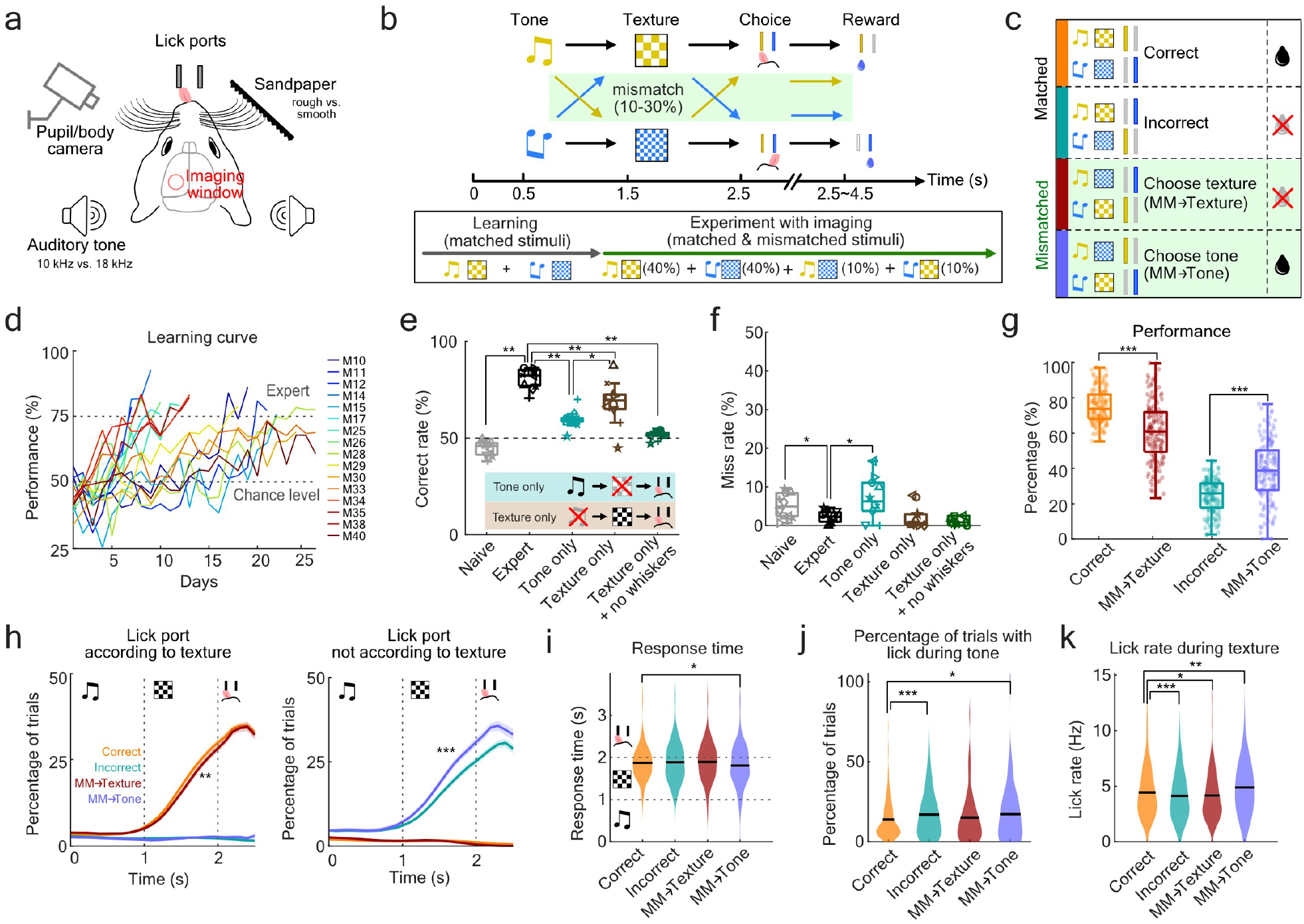
Mouse behavior in an auditory-cued texture discrimination task under matched and mismatched conditions. (**a**) Schematic of experiment setup with head-fixed mice with a 4-mm cranial window over S1 and PPC under a two-photon microscope. (**b**) Schematic of behavior paradigm. Mice were trained to discriminate two textures that were matched with two distinct auditory tone cues. After reaching expert level, tone-texture mismatch trials (10-30%) were introduced randomly within sessions. The choice window lasted maximally 2-s; it was terminated when the mouse licked during this window and the reward was delivered immediately. (**c**) Schematic, nomenclature and color code of different trial types. Trials were categorized based on the contingency of tone, texture, and choice. Colored bars on the left indicate the color code of each trial type. (**d**) Learning curves of all mice showing session-wise performance levels. M30, M33 and M40 were unstable performers: they did not reach expert performance level for entire sessions as shown here, but for multiple sub-sessions of 100 trials (see Supplemental Fig. 1a). (**e**) Performance in single-modality experiments, in which either only tone or only texture was presented. As a control, texture-only was also presented with whiskers trimmed. Markers indicate individual mice. The outlier (star) is M40 in (d). (**f**) Miss rates for experiments in (e). When only the tone was presented, mice were less engaged in the task. (**g**) Percentage of each trial type for matched and mismatched conditions. (**h**) Lick probability over trial time on the lick ports according to texture identity (left) or the opposite lick port (right), calculated as percentage of trials in each session with a lick event at each given time point. (**i**) Response time for different trial types. Mismatch-choose-tone condition shows a shorter response time overall (p = 0.0283). (**j**) Percentage of trials in each session, in which licks on the final choice spout were recorded during tone presentation. (**k**) Lick rate during texture presentation for different trial types. (e-f: n = 10 mice, 1 session per mouse, naïve was the average of first 3 sessions, expert was the average of best 3 sessions; g-k: n = 16 mice, total 148 sessions; Wilcoxon Signed-Rank test; here and in subsequent panels: *p < 0.05, **p < 0.01, ***p < 0.001)

During learning, the tone-texture pairing remained fixed to allow mice to gradually associate the tone with the upcoming matching texture (“matched trials”). Then, in expert mice, we randomly introduced 10-30% tone-texture mismatch trials (“mismatched trials”) to introduce a prediction error (Fig. 1b). In these trials, reward was given according to the tone, to encourage mice to generate active predictions. Under the conflict between tone and texture, mice could make two types of choices: when they chose the lick port according to the tone (“mismatch-choose-tone”, or “MM→Tone” trials), conceptually this situation corresponds to the case that the tone-based prediction, rather than the actual texture stimulus, dominated the decision; when mice chose according to the texture (“mismatch-choose-texture”, or “MM→Texture” trials), conceptually, the decision was made based on the actual texture rather than the tone-based prediction (Fig. 1c).

We trained 16 mice expressing GCaMP6f in L2/3 neurons (see Methods), all of which could successfully learn the task (Fig. 1d; see outliers in Supplemental Fig. 1a). We first analyzed the behavior of mice across task learning. Compared to naïve condition, expert mice showed suppressed licking during tone presentation, while the decision time shifted to the end of texture presentation (Supplemental Fig. 1b-e), indicating that mice associated the texture rather than the tone stimulus with reward. At the end of the experiment, we tested the performance of the mice upon presentation of either only tone or only texture while keeping the task structure unchanged. With these single-modality stimuli, mice could still perform the task above chance level, however at a reduced success rate. Furthermore, mice performed better for only-texture compared to only-tone presentation (Fig. 1e). The tone-only condition also resulted in a higher rate of task disengagement (miss rate, Fig. 1f), suggesting that mice regard the missing texture input as incomplete task structure. This also indicates that there was no significant re-learning of task rules from the mismatched trials, since these single modality experiments were done after all mismatched sessions. Finally, testing under texture-only condition with whiskers removed diminished task performance to 50% chance level (Fig. 1e). Together, these results indicate that mice rely on both tone and texture stimuli to perform the task, with texture being the most relevant stimulus, presumably due to its closer temporal link to trial outcome.

In mismatched trials, when an unexpected texture followed the tone, mice were less likely to decide according to the texture identity compared to matched stimuli (Fig. 1g, Correct vs. MM→Texture; in both these trial types the decision was made according to the texture identity). Similarly, tone biased the choice of mice to a degree that could not be explained by mere mistakes (Fig. 1g, Incorrect vs. MM→Tone; in both these trial types the decision was made opposite of the texture identity). When mice chose according to texture under mismatched condition, the lick probability during the texture and decision windows was slightly reduced (Fig. 1h left, Fig. 1k), indicating lower decision confidence. When mice chose according to tone, mice licked more decisively (Fig. 1h right, Fig. 1k), responded faster in general (Fig. 1i), and were more likely to lick before texture onset, during the tone window (Fig. 1j). It is worth noting that in the majority of trials, for all trial types, mice started licking after texture onset (Fig. 1h), indicating that the texture was the most relevant stimulus for the task. While there was no significant difference in the face and body movements across trial types (Supplemental Fig. 1f-h), the pupil diameter was higher across trial time when mice chose according to the texture under mismatched condition (Supplemental Fig. 1f,i), indicating a higher arousal state that could contribute to mice paying more attention to the texture input. Overall, these results indicate that, despite the prominence of the texture stimulus, the preceding auditory tone does bias the behavior and choice of mice.

### Simultaneous imaging of population activity in S1 and PPC areas during the task

Using a custom-built two-area two-photon microscope^32^, we simultaneously imaged the population activity in S1 and PPC when expert mice performed the task (Fig. 2a). As previously reported^26^, auditory and tactile stimuli recruit distinct PPC subdivisions: PPC-A (anterior) and PPC-RL (rostrolateral), respectively. To study interactions of cortical areas along the sensory hierarchy, we simultaneous recorded in two sets of experiments either S1 and PPC-A (9 mice, 40 sessions) or S1 and PPC-RL (14 mice, 91 sessions; 9 mice also had S1 and PPC-A sessions) (Fig. 2b). The location of S1 and PPC areas were determined by sensory mapping as well as retinotopic mapping^33^ (Supplemental Fig. 1a). We recorded in layer 2/3 across multiple depths (3-4 depths, between 100-300 µm), covering ∼50-600 neurons from each population (Supplemental Fig. 1b-c; Methods). Calcium indicator fluorescence signals from individual neurons and deconvolved spike rate were extracted using Suite2p^34^. All following analysis was performed on the deconvolved spike rate.

**Figure 2.**
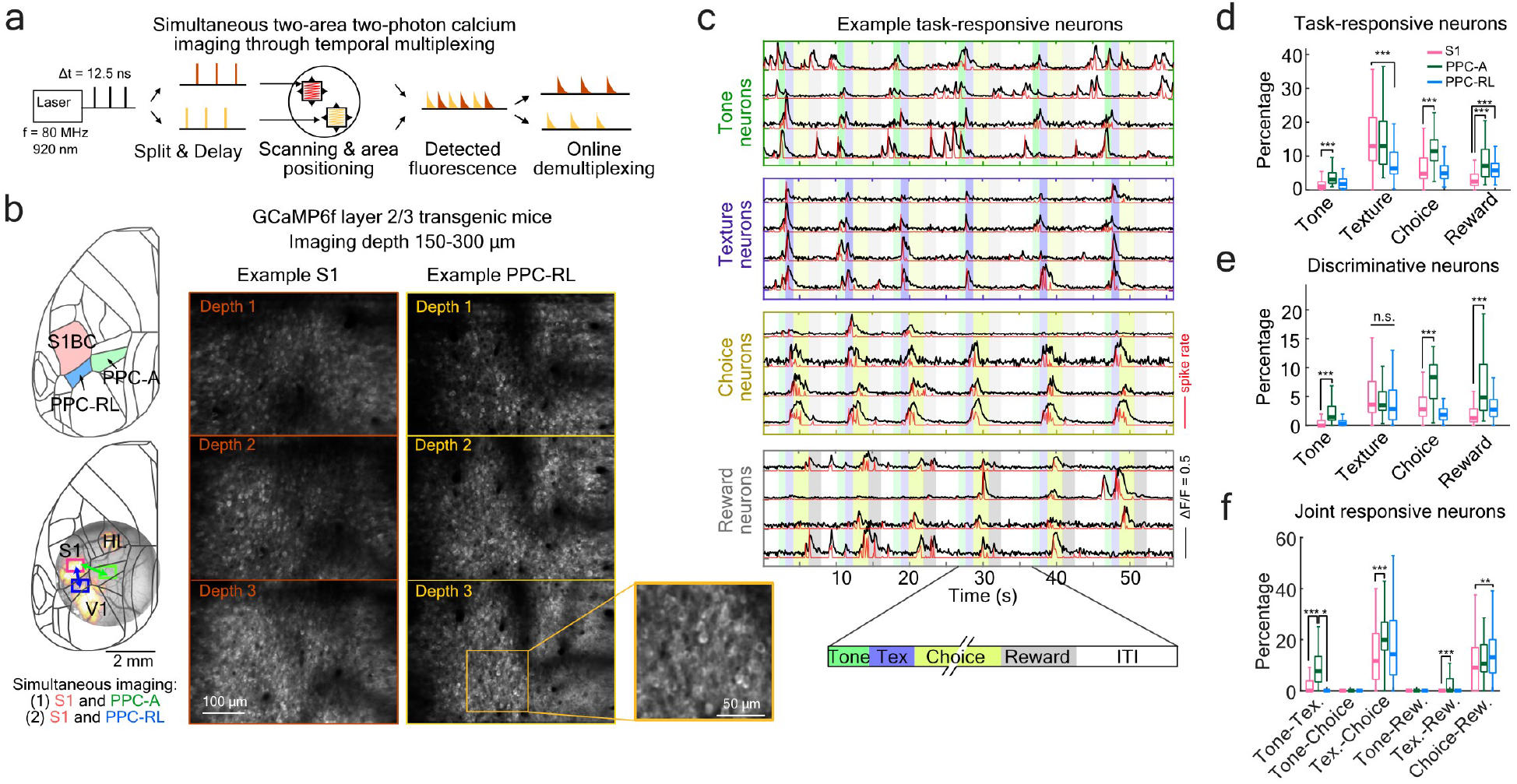
Simultaneous two-photon imaging of task-related S1 and PPC activity. (**a**) Temporal multiplexing-based simultaneous two-area two-photon imaging. Laser pulses were split into two copies, one of which was delayed by half of the pulse interval. Each copy was directed to an independently positioned field of view (FOV), and the emitted fluorescence was demultiplexed online with fast acquisition hardware. (**b**) Left: locations of S1, PPC-RL and PPC-A on the left hemisphere (top). Example widefield sensory mapping response, as well as example FOV locations, are shown in the bottom panel. Right: example FOVs of simultaneously imaged S1 and PPC-RL. (c) Example ΔF/F traces (black) and spike rate (red) of task-responsive neurons from one area in one imaging session. (**d**) Percentage of task-responsive neurons for each task window in S1, PPC-RL and PPC-A. (**e**) Percentage of discriminative neurons for each task variable in the three areas. (**f**) Percentage of joint responsive neurons (neurons that are responsive in two task windows). The percentage was calculated as N_i_⋂N_j_/N_i_⋃N_j_, where N_i_ and N_j_ are sets of responsive neurons for task phase *i* and *j*. (S1: 14 mice, 118 sessions; PPC-RL: 14 mice, 78 sessions; PPC-A: 9 mice, 40 sessions; Wilcoxon Rank Sum test).

All the three areas were engaged in the task, showing a varying degree of activation across the task windows. We identified task-responsive and discriminative neurons from matched conditions, using neuronal activity in each task window (Fig. 2c; Supplemental Fig. 2d; Methods). While S1 and both PPC-A and PPC-RL displayed highest task-related activity during the texture window, PPC-A had the highest fractions of both task-responsive and discriminative neurons in the remaining tone, choice, and reward windows (Fig. 2d-e), indicating its higher level position in the hierarchy. PPC-RL was less involved than PPC-A during sensory processing, particularly during tone presentation, but was also more engaged than S1 after reward delivery. Despite PPC-A being active in both tone and texture window, neurons that were active for both stimuli accounted only for a small percentage. Texture and choice, however, shared ∼20% of overlapping neurons (Fig. 2f). Overall, S1 was most tuned to texture processing, PPC-A was engaged throughout the trial time, and PPC-RL was involved in texture and reward processing.

### Tone-texture mismatch alters texture neuron tuning

We first asked whether tone-based prediction could alter the texture-evoked response of single neurons. To answer this question, we analyzed the response amplitude of tone- and texture-discriminative neurons. While tone-discriminative neurons showed differential responses to the distinct auditory tones, they did not show significant trial type-dependent activity during tone and texture presentation (Supplemental Fig. 3a-c). In contrast, texture-discriminative neurons activity could be altered by tone. In S1, texture-discriminative neurons showed strong texture selectivity only when mice chose according to texture under the mismatched condition; when mice chose according to tone, the activity of these neurons resembled the response to both the expected and the actual texture, showing mixed preferences (Fig. 3a-c, top panels). We obtained similar results in PPC (Fig. 3a-c, middle and bottom panels). In PPC-A, even more strikingly, the texture-tuned neurons showed strong preference to the expected rather than the actual texture when mice chose according to tone, but not when they chose according to texture (Fig. 3a-c, middle panels). This could not be explained by the joint choice selectivity of texture neurons, because we observed the same effect in PPC-A texture neurons that were not choice-selective (Supplemental Fig. 4a-b). Additionally, this inverted texture preference is distinct from the response distribution observed for incorrect trials, where mice had the same decision pattern (making the choice not according to the texture). These results provide strong evidence that tone-based prediction can shift the texture representation of individual neurons in PPC-A to the expected texture, while disrupting the texture representation in PPC-RL and S1.

**Figure 3.**
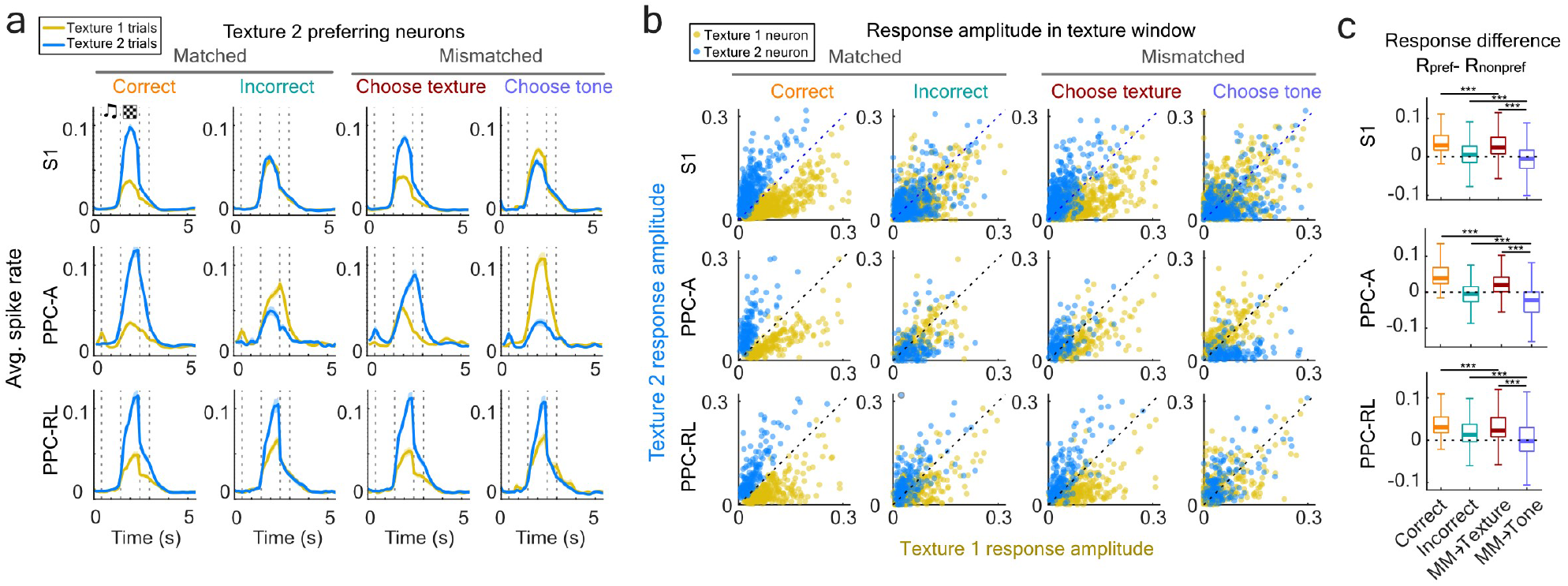
Tone-texture mismatch alters neuronal tuning to texture. (**a**) Average normalized spike rate of texture 2 discriminative neurons in S1, PPC-RL and PPC-A, in matched and mismatched trials. The spike rate of each neuron was normalized to be between 0 and 1 within each session. (**b**) Plots of texture 2 vs. texture 1 mean response amplitudes in the texture window for all texture discriminative neurons. Plots are shown across areas (rows) and trial types (columns). Each dot represents the responses of one neuron in one imaging session. (**c**) Selectivity index of the texture discriminative neurons during texture window, in different trial types. Selectivity index was calculated as the difference between the average response to the preferred texture and the average response to the non-preferred texture. (S1: 14 mice, 118 sessions, 1486 neurons; PPC-A: 9 mice, 40 sessions, 400 neurons; PPC-RL: 14 mice, 78 sessions, 618 neurons; Wilcoxon Rank-Sum test)

### Tone-texture mismatch alters population encoding

To further investigate the encoding of different task variables in the neuronal populations of these three areas and the representational changes, we trained linear support vector machine (SVM) decoders on the population activity patterns to discriminate tone (tone 1 vs. tone 2), texture (texture 1 vs. texture 2), choice (choose left vs. right), and reward (rewarded at left vs. right lick port) (Fig.4a-e; Supplemental Fig. 5a-b). Each decoder was trained and cross-validated using the time-concatenated spike rates of all neurons from the corresponding task window (e.g., texture window for the texture decoder; Fig. 4a), using trials with matched stimuli (excluding mismatched condition). Each trained classifier defines a hyperplane that best separates the two variables; by projecting the population firing rate onto the orthogonal axis of this hyperplane, or the “projection axis”, we could estimate the encoding strength of the corresponding task variable at each time point over trials^10^ (Fig. 4a; see Methods).

**Figure 4.**
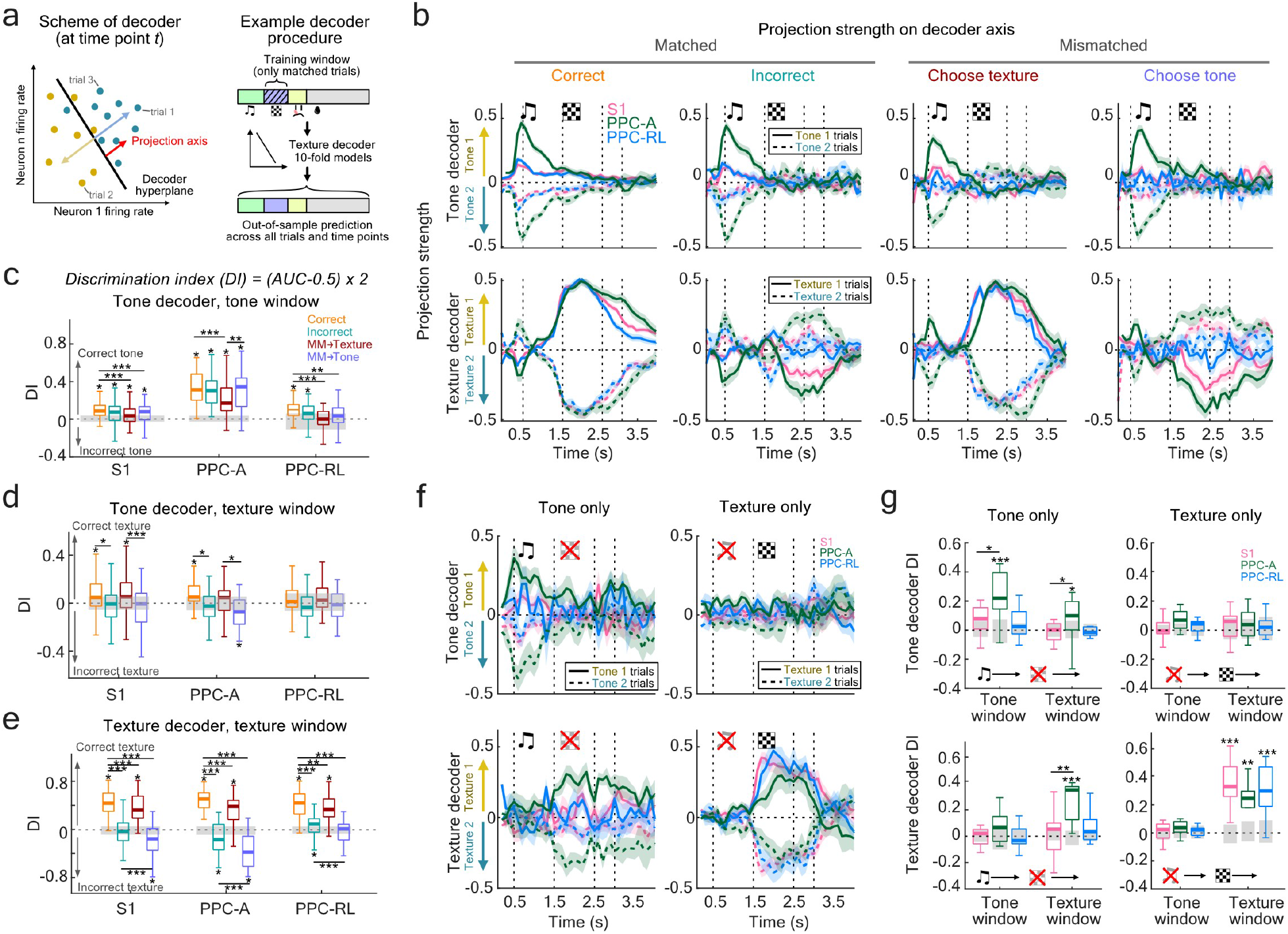
Tone-texture mismatch alters texture encoding in S1 and PPC populations. (**a**) Left: scheme of the decoders. Black line represents the decoder hyperplane; yellow and teal dots represent the neuronal population activity at a given time point in the trial; each dot was projected onto the orthogonal axis of the decoder hyperplane to represent the decoder confidence. Right: example procedure for texture decoder. (**b**) Neuronal population decoding of tone (top row) and texture (bottom row). Sensory encoding is represented by projection strength on the axes of linear decoders trained to discriminate tone and texture, in the tone and texture window, respectively. Line colors indicate area identity (magenta: S1; green: PPC-A; blue: PPC-RL); solid and dash lines indicate stimulus identity in the trial. (**c**) Discrimination index (DI) of tone decoder in the tone window. Stars above each box indicate significance compared to shuffled data (gray bars), for which neurons identities were shuffled, whereas trial and time correspondences were kept the same. Stars across boxes indicate comparison between trial types. (**d**) DI of tone decoder in texture window. (**e**) DI of texture decoder in texture window. (**f**) Decoder projection strength of tone (top row) and texture (bottom row) in single-modality experiments. (**g**) DI of tone decoder and texture decoder in single-modality experiments, in tone window and texture window, separately. Stars above each box indicate significance with shuffled data, stars across boxes indicate comparison between areas. (Wilcoxon Rank Sum test was used for comparison with shuffled data, Wilcoxon signed-rank test for comparison between trial types.)

Among the three areas, PPC-A best encoded tone information (Fig. 4b, top; Fig. 4c), consistent with the observation that PPC-A has the highest fraction of tone-tuned neurons and that it receives more input from auditory cortex than PPC-RL^26^. While the tone was mainly encoded in the tone window, its identity could still be read out after texture onset (Fig. 4d, gray bars indicate shuffled data). Interestingly, tone encoding strength corresponded to behavioral choice: in mismatch trials, when mice chose according to texture, PPC-A populations encoded the tone identity weaker as compared to correct trials (Fig. 4c); in contrast, when mice chose according to tone, tone encoding strength was comparable to correct trials. In the latter case, the tone identity (and thus the expected texture identity) was still decodable during texture presentation (Fig. 4d, middle). Similar results were observed in S1 and PPC-RL as well, despite weaker tone encoding in these two areas. These observations are consistent with the results that the tone could bias the behavior and choice of mice.

All three areas showed strong texture decoding performance (Fig. 4b, bottom). Consistent with the results in single neurons, in mismatched trials, the actual texture was encoded in all observed populations only when mice chose texture, although with less confidence compared to normal correct trials (Fig. 4e); when mice made their choice based on the tone, the expected texture instead of the actual texture was encoded (Fig. 4e). The timing of texture encoding is also worth noting. In correct trials under matched stimuli, the actual texture was encoded slightly before texture onset, indicating an active ongoing prediction (Fig. 4b, bottom left; Supplemental Fig. 5c). However, in mismatch-choose-texture trials, although the texture was encoded correctly, this early texture encoding was absent (Fig. 4b, bottom “Choose tex”; Supplemental Fig. 5c). This finding suggests a behavioral state of the mouse where it was waiting for and paying attention to the texture stimulus before texture onset (in agreement with the higher arousal state in Supplemental Fig. 1f,i). In addition, PPC-A texture encoding lagged behind of S1 and PPC-RL, suggesting that texture encoding was influenced by the preceding tone, which predicted the opposite texture identity, an effect particularly strong in PPC-A. On the other hand, in mismatch-choose-tone trials, the expected texture was encoded slightly before texture onset (Fig. 4b, bottom right; Supplemental Fig. 5c), and texture encoding in PPC-A preceded that of S1 and PPC-RL (Supplemental Fig. 5c), indicating that the tone facilitated the encoding of the expected texture. This effect was not due to choice-related activity, as choice could not be decoded during the tone window in these trials (Supplemental Fig. 5b, top). These results suggest that tone information in PPC-A led to encoding of the expected texture across S1 and PPC.

We next wondered whether the tone alone was sufficient to induce predictive texture encoding. To answer this question, we trained linear classifiers on trials where only the tone or only the texture was presented. Indeed, we found that tone stimulation alone resulted in strong texture encoding in PPC-A (Fig. 4f-g, left), and choice was only decodable in this area. In contrast, texture stimulation alone reduced the texture encoding speed and strength of PPC-A (Fig. 4f-g, right), but choice engaged both S1 and PPC (Supplemental Fig. 5d-e). This further suggests that PPC-A is able to generate predictive texture information based on the preceding tone, and it serves as a center for routing and transforming sensory information into decisions.

### Inter-areal interaction between S1 and PPC during task trials

We next asked if such predictive information could be explained by the top-down and bottom-up interactions between S1 and PPC areas during the task. We started by characterizing the interaction structure of S1 and PPC populations during the task. To measure the interaction between neuronal populations, we used canonical correlation analysis (CCA) due to its symmetric way of treating the two populations. CCA has been applied to analyze the inter-areal interactions in several recent studies, for both electrophysiology and calcium imaging data^16,17,19^. In the framework of CCA, the activity of a neuronal population can be represented in a high-dimensional space, where each dimension represents the activity of one neuron in this population. The population activity at a given time point in a given trial is represented as a point in this space. With two simultaneously recorded populations, CCA finds pairs of dimensions that maximize the correlation between the projections of two populations (Fig. 5a). These pairs of dimensions (canonical dimensions) define the shared interaction axes of the two populations at this time point.

**Figure 5.**
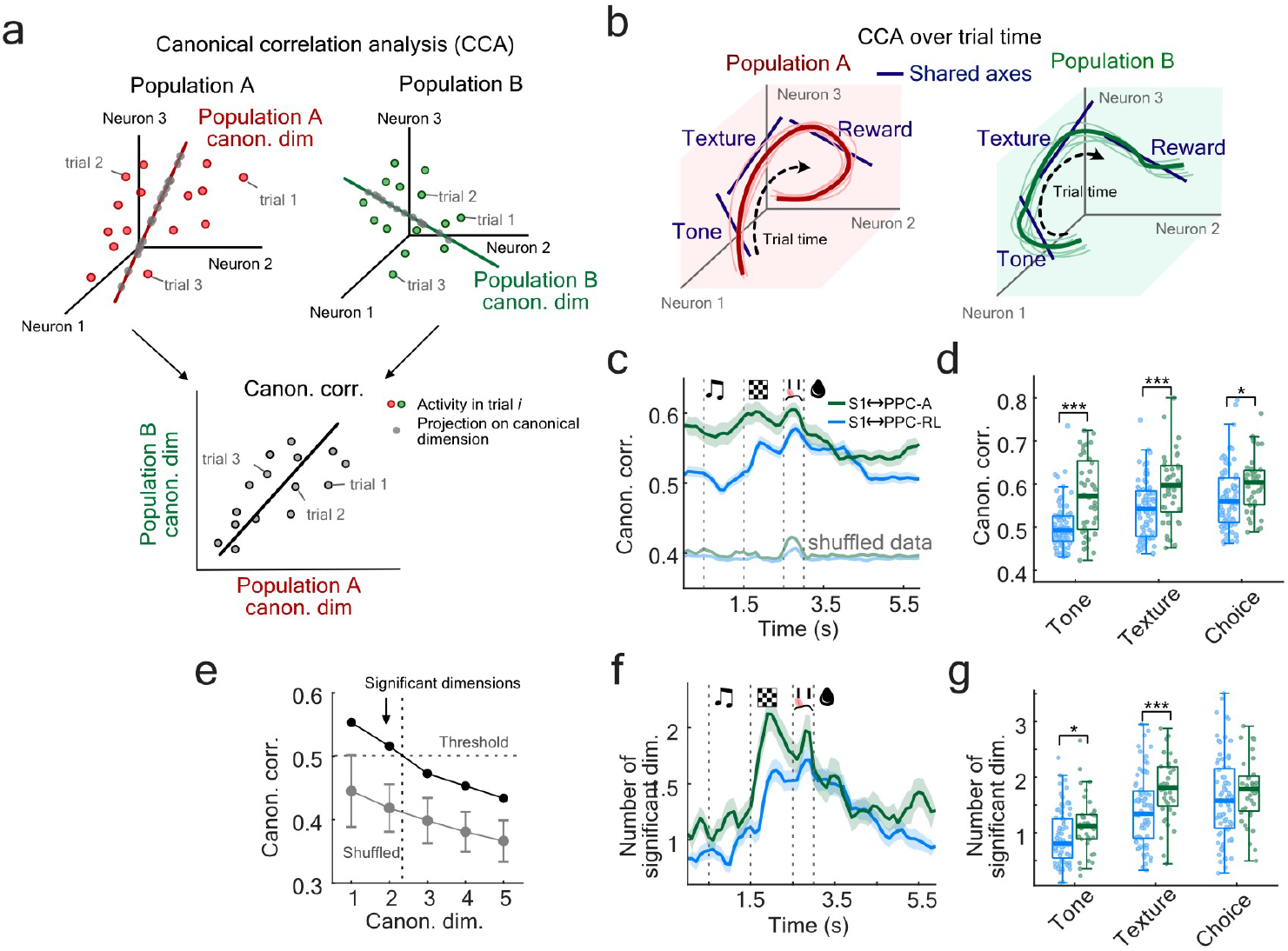
Interaction pattern of S1 and PPC areas during the behavioral task. (**a**) Illustration of canonical correlation analysis (CCA). The activity of each neuronal population can be represented as points in a high-dimensional space, where each dimension represents the activity of one neuron in this population. At a given time point in the trial, CCA identifies a set of canonical dimensions through linear combinations of variables from the two populations that have maximum correlation. (**b**) Illustration of defining separate CCA axes over the trial time. (**c**) Top canonical correlation between S1 and PPC-A (green), and S1 and PPC-RL (blue). Light colors indicate shuffled correlation, where only trial correspondence between the two areas were shuffled, while trial structure was kept the same. (**d**) S1 and PPC-A interaction was consistently higher than S1 and PPC-RL interaction across trial time. (**e**) Number of significant interaction dimensions were determined by generating shuffled correlations and defining a threshold (mean + 3 S.D.) using the first correlation values. Canonical dimensions in real data with correlations higher than this threshold were considered significant. Example is from one imaging session. (**f**) Number of significant dimensions in S1 and PPC-A, and S1 and PPC-RL pairs. (**g**) The number of significant dimensions for S1 and PPC-A was also higher than S1 and PPC-RL across trial time. (S1↔PPC-A: 9 mice, 40 sessions; S1↔PPC-RL: 13 mice, 71 sessions; Wilcoxon Rank Sum test)

Stable CCA models depend on having sufficient amount of data (time points) compared to the number of variables (neurons)^35^. For a typical dataset, we have ∼200 neurons per area, and ∼300 trials per session,which is rather challenging for forming a stable CCA solution. To ensure that the discovered interaction dimensions are stable, we first applied principal component analysis (PCA) to both populations and kept the first 30 components. As we are interested in discovering the interaction between the two populations, we then subtracted the stimulus-triggered average to keep only the residual activity that captures trial-to-trial co-variation. Interaction between brain areas is not static^16,17^; to capture such changing interaction, we then applied CCA on the residual activity of the two populations using a sliding 0.5-s window across the trial time (Fig. 5b). For the best-correlated interaction axis (the first canonical dimension), S1 and PPC-A showed consistently stronger interaction over trials than S1 and PPC-RL (Fig. 5c-d). S1 and PPC-A interaction was the lowest during the inter-trial interval, and ramped up during tone presentation, while S1 and PPC-RL interaction only increased after texture onset. We then wondered whether the number of communication subspace between S1 and PPC changes during trial time. To answer this question, we determined the number of significant canonical dimensions, defined as the dimensions with canonical correlation exceeding the highest shuffled correlation, where the trial correspondence between the two populations was randomized (Fig. 5e). Consistent with previous reports^15,16^, S1 and PPC interacted in low-dimensional space (1-2 dimensions), with texture and choice involving higher interaction dimensions in both pairs of areas (Fig. 5f). During tone and texture window, S1 and PPC-A interacted in higher-dimensional subspaces than S1 and PPC-RL (Fig. 5g), indicating PPC-A was more involved in sensory processing during the task. In the following analysis, we focused on the first interaction dimension with the highest correlation.

### Top-down and bottom-up interactions govern behavioral choice

In behavioral tasks, sensory information flows through the cortical hierarchy to generate decisions. For our task, a particularly interesting question is whether the behavioral choice of mice under mismatch condition could be explained by the net output of top-down (prediction) and bottom-up (sensory) information. Conceptually, choosing texture could be explained by stronger bottom-up texture information in S1 than top-down tone-based prediction in PPC, while choosing tone could be explained by stronger top-down prediction than bottom-up sensory information. To test this hypothesis, we measured the strength of bottom-up and top-down information flow by introducing a temporal lag in the CCA models (Fig. 6a). We defined bottom-up strength by moving S1 activity window ahead of PPC and averaging the top canonical correlation across lags (averaged over 0.3 s), and top-down strength by moving PPC activity ahead of S1. The information flow is represented as the net effect (the difference between bottom-up and top-down strength) normalized by the total interaction strength (the sum of bottom-up and top-down strength) (Fig. 6a).

**Figure 6.**
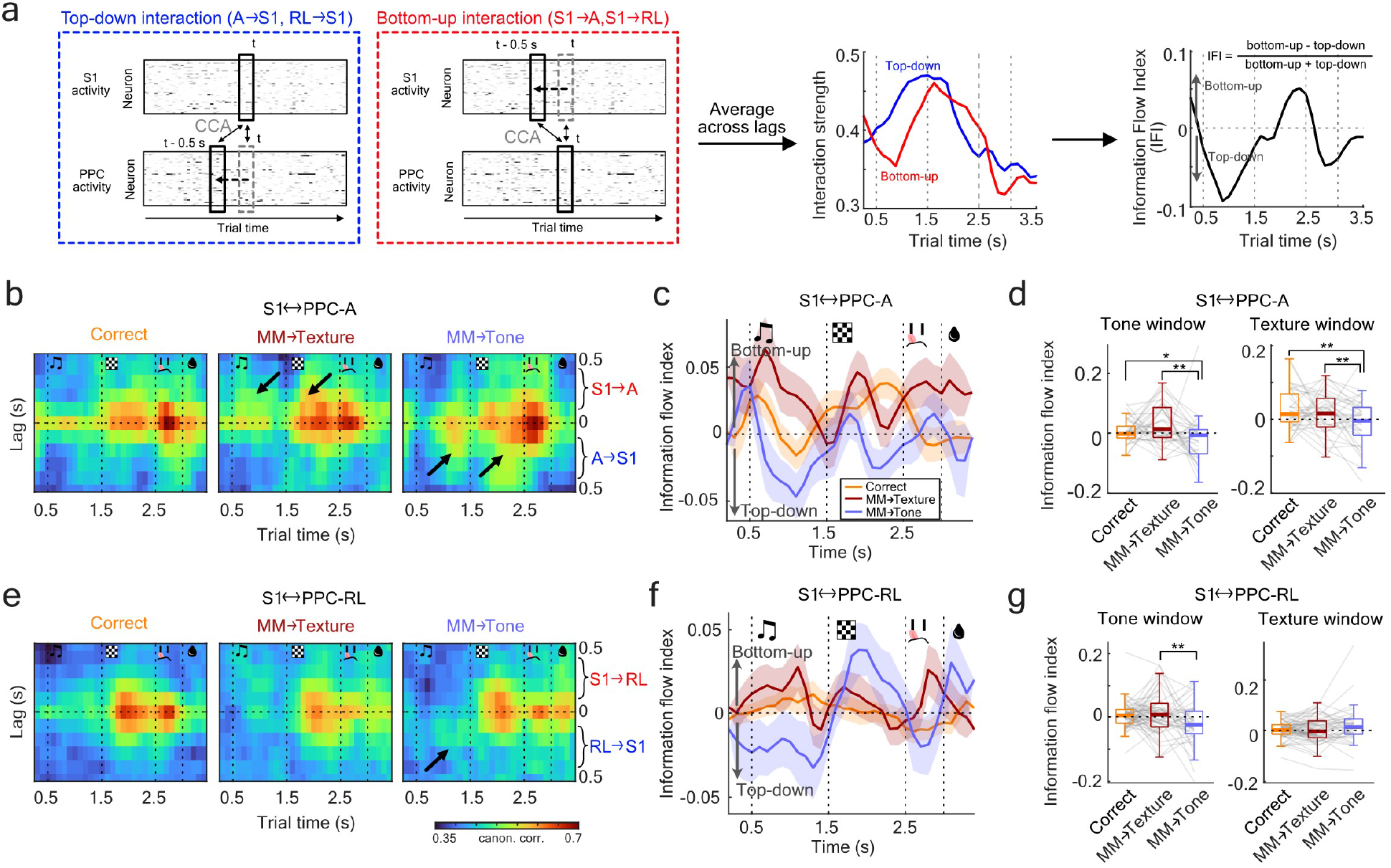
Top-down and bottom-up interactions between S1 and PPC areas during prediction mismatches. (**a**) Top-down interaction strength (PPC-A/RL to S1) was calculated by the average canonical correlation with a negative lag for PPC, and bottom-up interaction strength was calculated by the average correlation with a negative lag for S1 (left and middle panels). The direction and strength of S1-PPC interaction was characterized by the information flow index (IFI), defined as ratio of the difference between the two interactions to their sum (right panel). In this example dataset, the population interaction was first top-down dominant, then switched to bottom-up dominant before top-down took over again. (**b**) Lagged canonical correlation between S1 and PPC-A, averaged across all sessions, for different trial types. (**c**) Information flow index quantified from (b). (**d**) Quantification of information flow index between S1 and PPC-A. (**e**) Lagged canonical correlation between S1 and PPC-RL, averaged across all sessions, for different trial types. (**f**) Information flow index quantified from (e). (**g**) Quantification of information flow index between S1 and PPC-RL. (Mice and session numbers are the same as Fig. 5; one-sided Wilcoxon Signed-Rank test)

In matched trials, S1 and PPC-A interaction showed strong bottom-up dominance during texture presentation (Fig. 6b-c, correct). In mismatched trials, when mice chose texture, both tone and texture windows were dominated by bottom-up information (Fig. 6b-d), with weaker tone information and stronger texture information. This indicates that mice could be in an internal state that attended more to texture input. Since mice mostly relied on texture for solving the task (Fig. 1e-f, MM→Texture; Supplemental Fig. 1b-e), this result is in agreement with the higher arousal state observed in these trials (Supplemental Fig. 1f,i). In contrast, when mice chose tone, tone presentation induced strong top-down dominance from PPC-A to S1, and during texture presentation, such top-down influence persisted (Fig. 6b-d, MM→Tone). This top-down influence was also observed from PPC-RL to S1 during tone presentation (Fig. 6e-g), indicating an internal state that emphasized top-down inputs.

The task-related information flow between S1 and PPC was in agreement with the results from single-modality experiments. When mice were presented only with the tone, the population interaction resembled that observed for the mismatch-choose-tone condition: During the tone, a strong top-down information flow occurred from PPC-A to S1 and from PPC-RL to S1 (Supplemental Fig. 6a-f, tone-only condition), and such top-down flow persisted throughout the texture window (where no texture was presented) between PPC-A and S1, consistent with role of PPC-A in tone-based predictions. When only the texture was presented, bottom-up flow from S1 to PPC-A during the texture stimulation was observed instead (Supplemental Fig. 6a-f, texture-only condition), resembling the interaction observed for the mismatch-choose-texture condition.

To exclude the possibility that these observations were specific to the CCA method we used, we additionally computed population interaction strength with Pearson correlation as an alternative approach (Supplemental Fig. 7a). Overall, Pearson correlation reproduced all the results, even though Pearson correlation does not capture the optimal interaction between populations: we again observed higher interaction strengths between S1 and PPC-A, particularly during the texture window (Supplemental Fig. 7b-c), and strong top-down influence from PPC-A to S1 in mismatch-choose-tone trials, in both tone and texture window (Supplemental Fig. 7d-e). A top-down dominance during tone was also observed in PPC-RL and S1 (Supplemental Fig. 7f-g). When only the tone was presented, both pairs of areas showed similar interaction patterns as mismatch-choose-tone trials, whereas the texture-only condition featured stronger bottom-up inputs (Supplemental Fig. 7h-k). Together, these results demonstrate how the dynamic top-down and bottom-up interactions between S1 and PPC, particularly PPC-A, govern the behavioral choices of mice during the task (Fig. 7a-c).

**Figure 7.**
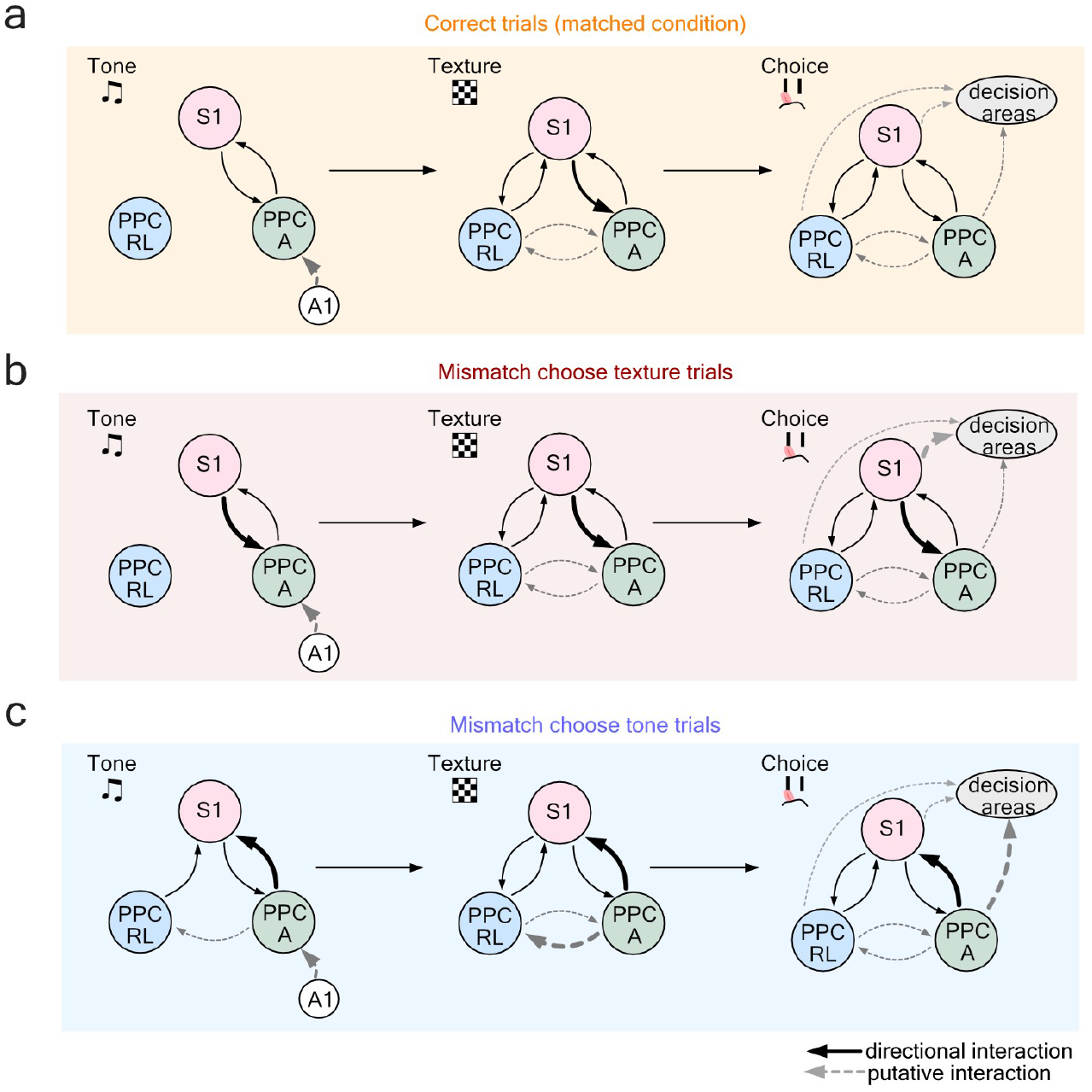
A model of S1, PPC-A, and PPC-RL interactions during the behavioral task. (**a**) In the correct trials, tone information is encoded in PPC-A, potentially through inputs from auditory cortex (A1). PPC-A further generates predictions about the upcoming texture and communicates with S1. During the texture presentation, S1 sends texture information back to PPC-A, while all three areas are involved in the processing and transformation of texture information. The decision is formed through coordinated efforts from S1 and PPC. (**b**) In mismatch-choose-texture trials, mice are in a state that attends more to S1 inputs. During texture presentation, strong texture information in S1 is sent bottom-up to PPC-A that eventually leads to the behavioral choice. (**c**) In mismatch-choose-tone trials, tone presentation generates stronger tone encoding and texture prediction in PPC-A, which is sent to S1 during the tone and texture windows, overwriting the bottom-up texture input from S1. Thus, the choice is made according to the predicted texture. Solid arrows represent directional interactions between recorded areas, dashed arrows represent putative interactions between pairs of areas that were not recorded simultaneously in this study.

## Discussion

In this study, we utilized a two-area two-photon microscope to study the interaction between the sensory S1 area and the next-higher association area PPC during an auditory-cued texture discrimination task. We focused on cortico-cortical interactions related to the predictive processing theory, by introducing tone and texture mismatches to induce conflicts between tone-based top-down texture predictions and bottom-up tactile input. We demonstrated that when predictions dominated sensory inputs, as in trials in which mice made their decisions based on tone instead of actual texture identity, both single-cell and population encoding of texture were disrupted in S1, while PPC-A encoded the expected texture. This situation also corresponded to a stronger top-down information flow from PPC-A to S1. When sensory input overrode predictions and mice decided based on the actual texture identity instead of the predicted one, texture encoding in S1 and PPC remained unchanged, and bottom-up information flow from S1 to PPC-A was stronger (summarized in Fig. 7a-c). Overall, our results provide strong evidence for a cortical implementation of predictive processing in the context of sensory-driven decision making.

Although predictive processing provides an attractive framework for understanding a wide range of brain functions, the specific neural circuits involved and the neuronal population mechanisms are still poorly understood. It has been shown that primary sensory cortices can develop experience-dependent predictions that impact stimulus response: expected stimuli are suppressed, and unexpected stimuli are amplified^6,7,36,37^. This occurs for both unimodal stimulation where repetitive stimuli are presented to the animals^7,36,37^, and multi-modal stimulation where animals associate two stimuli (e.g., auditory and visual)^6,13^. In addition, expected stimuli are encoded faster in the gustatory cortex when predicted by a leading auditory tone^8^, providing another benefit of predictive processing in sensory cortices. Such predictions can be conveyed directly between primary sensory cortices^6^, but also depend on long-range projections from higher areas^7,14,38^. While frontal areas such as the anterior cingulate cortex have been more extensively studied in predictive processing, the role of posterior association areas in this process is not well understood. Here, we provided evidence for a missing step in this pathway: the association area PPC-A carried sufficient predictive information based on a tone to encode the expected texture and choice even in the absence of texture. Thus, PPC-A, together with S1, forms part of the hierarchical cortical circuit for sensory-based predictive processing for this cross-modal behavioral task.

How cortical areas communicate with each other and what information is exchanged between them during active behavior, are still open questions. It has been shown that the direct projections from higher cortical areas to primary sensory areas carry specific task-related information such as expected stimulus and reward^7,9,14^, providing one mechanism for how top-down predictions affect the response of primary sensory areas. However, due to the technical challenges of simultaneously recording from sufficient numbers of neurons in multiple areas, studies concerning the interactions between neuronal populations across cortical areas are still scarce. Recent studies have demonstrated that interactions between primary and higher areas usually occur in low-dimensional communication subspaces that do not necessarily align with the stimulus-encoding subspace in each area^15,39^. In addition, interactions between areas are dynamic and vary throughout stimulus presentation and behavioral tasks^16–19^. Our study adds to these findings by providing a detailed analysis of the dynamic population interactions that occur during a cross-modal sensory discrimination task. Our results suggest that the task-relevant variables are encoded in low-dimensional latent spaces in S1 and PPC and that the dominance of either top-down or bottom-up processing corresponds to perceptual choices during the task, putting more weight on either predictions or actual sensory inputs. These findings are consistent with the predictive processing theory and contribute to our understanding of the mechanisms underlying cortical information transformation and communication.

Specifically, we observed a disrupted texture encoding in S1 accompanied by stronger top-down information flow, indicating that top-down predictions are able to change primary sensory cortex responses. The specific multi-modal task design in this study contributed to the sensory prediction, because PPC is involved in multisensory processing^22,23,25,40^ and therefore is well-positioned to develop cross-modal predictions. In PPC as well as primary areas, inhibitory interneurons play a key role in suppressing expected stimuli and augmenting unexpected stimuli^23,41,42^. Specifically, disinhibition through top-down projection from PPC to S1 provides a possible local mechanism for our observation of altered sensory encoding in S1. Furthermore, other brain areas such as the thalamus are likely to be involved in the relaying of prediction as well: higher-order thalamic nuclei are interconnected with many cortical areas^20,43^, providing another information routing station for processing of sensory conflicts by comparing top-down and bottom-up information.

The composition and exact locations of rodent PPC and its subregions are still a topic of debate^22,26,44^.

Historically, rodent PPC has been described as a part of higher visual areas, with PPC-RL largely overlapping with VISrl, and PPC-A overlapping with VISam^33,44^. However, PPC not only receives visual input, but also extensive inputs from somatosensory, auditory and olfactory cortices, as well as higher cortical areas^22,45^. Most PPC studies so far have defined PPC as a single area with varying coordinates, leading to contradictory results in some cases^23,46–48^. In our study, the two subregions of PPC, PPC-A and PPC-RL, exhibited different task relevance. While PPC-A was involved in all stages of the trial, PPC-RL was only involved in texture and choice processing. This is consistent with the reports that PPC can be divided into modality-specific subregions: the medial PPC-A more auditory, while the lateral PPC-RL is more tactile^26,45^. The inputs from the auditory cortex and the somatosensory cortex to PPC seem to form a gradient along the medial-lateral axis, suggesting a functional continuity of PPC^45^. In addition, these inputs presumably interact with and integrate inputs from primary visual cortex, which can be relevant in other cross-modal behavioral tasks. PPC is not only a center for multisensory processing, but also participates in decision making ^29,46,49,50^, via its dense connectivity with frontal areas^22^. Therefore, subregions in PPC likely serve as a continuum of intermediate routing stations of weighted sensory information flow towards frontal areas, and top-down information back to sensory cortices. Further work is required to characterize the specific roles of PPC subregions and their interactions in more details.

### Online Methods

All procedures of animal experimentation were carried out according to the guidelines of the Veterinary Office of Switzerland and following approval by the Cantonal Veterinary Office in Zurich (licenses 234/2018, 211/2018).

### Mice and dataset

16 mice were included in this study. Mice were one of the following strains: RasGRF2a-dCre;CamK2a-tTA;TITL-GCaMP6f (M10, M11, M12, M25, M26, M28, M29, M33, M34, M35), GP5.17(C57BL/6J-Tg(Thy1-GCaMP6f)GP5.17Dkim/J, Jackson Laboratory 025393) (M14, M15), Snap25-IRES2-Cre-D;CamK2a-tTA;TITL-GCaMP6f (M17), RasGRF2a-dCre;tTA2-GCaMP6f (M30, M38, M40). All transgenic strains express GCaMP6f in layer 2/3 pyramidal neurons of the neocortex. Both sexes were included in this study (male: M10, M14, M15, M25, M26, M30, M33, M34, M35; female: M11, M12, M13, M17, M28, M29, M38, M40). All mice were adults (12-16 weeks old) when experiment started. Out of the 16 mice, S1-PPC_A_ imaging was performed on 9 mice (M25, M26, M28, M29, M30, M33, M34, M35, M40); S1-PPC_RL_ imaging was performed on 14 mice (M10, M11, M12, M14, M17, M25, M26, M28, M29, M30, M33, M34, M35, M40). Two mice (M15, M38) were only included in behavioral studies but not in the neuronal data analysis due to the decayed cranial window quality. One mouse (M17) was removed from cross-area analysis due to decayed S1 imaging quality from lateral skull bone growth under the cranial window.

### Surgical procedures

A craniotomy was performed on all mice over S1 and PPC in the left hemisphere. Mice were anesthetized with 2% isoflurane mixed with oxygen, and body temperature was maintained at 37℃. After analgesia treatment (Metacam, 5 mg/kg, s.c.; lidocaine gel over the skull skin) and locally, the skull was exposed, a 4 mm round cranial window was made with dental drill, and covered with glass coverslip using dental cement (Tetric EvoFlow). A light-weighted head-bar was fixed on the skull using dental cement. After the surgery, animals were continually monitored for three days, and treated with analgesics (Metacam, 5 mg/kg, s.c.). For strains that expressed destabilized Cre (dCre), we induced stable GCaMP6f expression by administering trimethoprim (TMP, Sigma T7883). TMP was reconstituted in Dimethyl sulfoxide (DMSO, Sigma 34869) at a saturation level of 100 mg/ml, and intraperitoneally injected (150 mg TMP/g body weight; 29 g needle) at least one week before imaging commenced.

### Behavior training

All mice were kept under a reversed dark-light cycle. Mice were allowed to recover for at least 1 week before behavior training started. Before behavior training, mice were first accustomed to the hands of the experimenter at the home cage for several days until showing no sign of stress, then accustomed to head fixation. Then, mice were put under water scheduling, and were introduced to the behavior setup. During the first 2-3 sessions, mice were given sugar water reward from one of the two lick ports after the choice tone (2 beeps at 3 kHz of 50-ms duration with 50-ms interval). Once they learned to lick after the choice tone to obtain the water reward, we introduced the textures. Textures were followed by reward delivery on the corresponding lick port upon licking, and licks on the wrong port did not result in punishment or trial abort. This stage lasted for 1-2 sessions. Once mice were accustomed to the trial structure, we started formal training.

Behavior training was carried out using a custom written LabView software. Each trial started with one of two distinct auditory tones (10 kHz or 18 kHz, 6 repetitions, 50-ms duration and 50-ms intervals). One second after tone onset, the presentation of one of two distinct textures followed (sandpaper, P100 vs. P1200 for M10-17, P280 vs. P800 for M25-40). A rotary motor “swung” the texture onto the whisker pad from the top. The texture was presented for 1 s and then moved away from the mouse with a linear motor stage. At the end of the texture window, choice window started, indicated by the choice tone described above. The choice window lasted for up to 2 s. As soon as mice licked during the choice window, it was terminated and the reward window started. If mice chose the correct lick port, a small water reward was delivered (∼4 µl sugar water); wrong choices were not punished. The inter-trial interval was randomly distributed between 4-8 seconds.

During training, when a mouse made incorrect choices, the same tone-texture stimulus pair was presented again in the following trial until the mouse chose correctly. This “repeat incorrect” strategy facilitates learning and prevents the mice from forming a bias towards one of the two lick ports. If the mouse disengages from licking, in 10% of these miss trials the reward was delivered after the choice window to motivate the mouse. Each day, the training lasted as long as the mouse was actively engaged in the task, typically 200-400 trials. Training was done once per day, 5-6 days per week. Weight, health, and water intake were monitored daily. All training was performed in the dark and monitored through a behavior camera with a small infrared light source. Mice were considered experts when they reached 75% correct performance for three sessions. Among all mice, three mice showed an unstable behavior, with relatively high fluctuations of the within-session performance (Supplemental Fig. 1a); however, they all showed clear signs of learning, reaching sub-session performance peaks above expert level for consecutive days. We attributed the unstable behavior to environmental stress; in particular, the training of M40 coincided with construction work in the animal facility, resulting in longer training duration.

### Mismatched trial design

After mice became stable experts, we started introducing mismatched trials. In these sessions, the first 20-30 trials were with matched stimuli (without mismatch), and only afterwards mismatch trials were randomly presented in 10%-30% of the trials. To avoid confusion for mice and prevent re-learning of new rules, we kept the repeat incorrect strategy throughout these sessions for matched trials, reinforcing the learnt rules. Because the task is mostly a texture-dependent task (see Results section, Fig. 1, and Supplemental Fig. 1), we rewarded according to the tone in mismatched trials, in order to encourage mice to pay attention to the tone and generate more mismatch-choose-tone response.

### Single sensory modality experiments

Tone-only and texture-only sessions were done at the end of the experiment, after mice completed all mismatched sessions (7-12 sessions). In tone-only sessions, trials started with a 1-s auditory tone, as for the matched pairing condition. Afterwards, the rotary motor carrying the texture swung in, generating the same motor noise, but stopped above the whisker pad of mice. Therefore, texture was “presented” above the mice, out of reach for their whiskers. Choice window, reward window and inter-trial interval were the same as in matched condition. In texture-only sessions, the tone before the texture presentation was omitted. Trial started with a 1-s window with no sensory stimulus, followed by normal texture presentation, and then choice and reward windows. These single modality sessions typically were restricted to 100-150 trials to prevent re-learning.

After the single-modality experiments, we trimmed the whiskers of the mice, and conducted another texture-only session. This session served as a control experiment to exclude the possibility that mice relied on other environmental cues (visual, olfactory, etc.) to perform the task.

### Behavior monitoring

Face and body movements as well as the pupil diameter of the mice were monitored and recorded using a CMOS infrared-sensitive camera (Basler acA1440-220um). A small 940-nm infrared LED was positioned in front of the mice to illuminate their face and body. Since mice were in complete darkness, their pupils were dilated by default. To monitor the pupil diameter with a larger dynamic range, we restrained the pupil by carefully positioning a small UV LED (385 nm, Thorlabs LED385L) close to the eye contralateral to the texture presentation. Trial-related behavior was recorded at 50 Hz, simultaneously with calcium imaging, triggered by each trial start. Licking was recorded throughout tone, texture, and choice windows, and was estimated based on the event rate from the capacitive lick sensor sampled at 100 Hz.

To extract body and face movements, we manually selected two regions of interest (ROIs), one on the whisker pad, the other on the forelimb and chest region. Movement was calculated as frame-to-frame variation by computing (1-corr(*f*_t_, *f*_t+1_)), where corr(*f*_t_, *f*_t+1_) denotes the frame-to-frame correlation of the ROI. We tracked the pupil diameter using a custom MATLAB script: we first manually selected an ROI over the eye region, then binarized the pupil (pupil was bright due to two-photon illumination of the cortex at 920 nm). Then, pupil diameter was estimated by fitting the binary region to an ellipse. Body and face movements, as well as pupil diameter, were smoothed with a median filter of 200-ms width.

### Sensory mapping

To determine the exact locations of S1 and PPC, we performed widefield sensory mapping on all the mice before the two-photon imaging sessions started, following a previously described procedure^26^. Mice were lightly anesthetized under 1% isoflurane and kept at 37℃. Three types of sensory stimuli were delivered: visual, whisker, and hindlimb. For visual stimulation, a small blue LED was positioned close to the eye contralateral to the cranial window, and a brief 200-ms flash was presented, followed by 10-s recovery time. For whisker and hindlimb stimulation, a loud speaker-coupled vibrating bar was used to induce a vibrating touches of the whiskers and hindlimb paw (20 Hz for 2 s) on the side contralateral to the cranial window. Each stimulus modality was repeated 30 times.

Widefield imaging was simultaneously performed through the cranial window. A blue LED light source (Thorlabs; M470L3) was used for excitation of GCaMP6f, together with an excitation filter (480/40 nm BrightLine HC). The excitation light was directed to the cranial window through a tube lens (Thorlabs TTL100-A) followed by a 4x objective (Thorlabs TL4X-SAP, NA 0.2). A dichroic mirror (510 nm; AHF; Beamsplitter T510LPXRXT) was positioned between the two objectives to separate excitation and emission light. Emission light was filtered (emission filter 470/10 nm, Edmund Optics) and recorded with a sensitive CMOS camera (Hamamatsu Orca Flash 4.0).

To obtain a more precise location of PPC-RL and PPC-A, we further generated a retinotopic map through visual field sign mapping, following a previously described procedure^33^. Briefly, a drifting spherically-corrected checkerboard visual stimulus was presented on an LED screen (Adafruit Qualia 9.7” DisplayPort Monitor, 2048×1536 Resolution) across the visual field of the mice at 0.043–0.048 Hz. The stimulus sequence consisted of four cardinal directions, each presented with 10 repetitions. The screen was positioned in front of the eye contralateral to the cranial window, such that the stimulus covered retinotopic locations from approximately -20 to +30 degrees in altitude and -10 to +90 degrees in azimuth. The retinotopic map was calculated using previously reported analysis pipeline^33^. The final location of S1, PPC-A and PPC-RL was determined by optimally aligning the sensory map and retinotopic map together to the Allen Mouse Common Coordinate^43^.

### Two-area two-photon imaging

Two-area two-photon imaging was performed using a custom-built microscope that has been previously reported^32^. The simultaneous two-area imaging was implemented through a temporal multiplexing technique, where the laser pulse train from a Ti:sapphire laser (Mai Tai HP DeepSee, Spectra-Physics) was split in two temporally interleaved copies, each directed through an independently movable unit to a separate field of view. Each beam path was equipped with an electrically tunable lens (Optotune EL-10-30-C) to enable rapid focal changes for imaging multiple depths. Imaging was done at 920-nm excitation with a green emission filter (510/42 nm bandpass). A 16x objective was used (N16XLWD, Nikon, NA 0.8). The microscope was controlled by a custom-written software Scope (http://sourceforge.net). Calcium imaging was acquired at 3 different depths in layer 2/3, separated by 40-50 µm. The FOV size was ∼450×500 µm, at a resolution of ∼370×256 pixels. The volume rate was typically ∼9.32 Hz for two areas and three imaging depths per area. Laser power was adjusted for each plane, at 40-70 mW under the objective. For each mouse, the FOV positions and/or depths were slightly adjusted for each imaging session to cover different neuronal populations in S1 and PPC. Imaging was triggered by the start of each trial, and the image acquisition finished 0.5 s before the end of the inter-trial interval (i.e., the start of the next trial). During the duration of the experiment, most mice maintained a clear window with good imaging quality. The number of imaging sessions for each mouse is as following: For S1-PPC_A_ imaging, the number of sessions was 3, 4, 5, 5, 4, 4, 7, 4, 4, for M25, M26, M28, M29, M30, M33, M34, M35, M40, respectively. For S1-PPC_RL_ imaging, the number of sessions was 10, 9, 10, 2, 1, 8, 5, 4, 8, 5, 3, 4, 4, 3, for M10, M11, M12, M14, M17, M25, M26, M28, M29, M30, M33, M34, M35,M40, respectively. Note that the number of PPC-RL sessions was higher for M10, M11 and M12 due to a lower percentage of mismatch trials (∼10% mismatch trials).

### Processing of two-photon imaging data

We used Suite2p to extract neuronal traces^34^. This pipeline includes a rigid motion correction on the raw data, a model-based background subtraction, a neuron-identification algorithm, fluorescence extraction, and a neuron classifier. Raw fluorescence and neuropil traces were extracted from identified neurons, and neuropil-corrected traces were obtained. A deconvolution algorithm was applied to the corrected fluorescence traces to estimate the spike rate of neurons (in arbitrary units). Spike rate was further normalized by the baseline F0 estimated by Suite2p. We tuned the neuron classifier in the above pipeline to identify potential neurons, and we further manually curated each dataset to discard non-neuronal structures or low-quality ROIs. All analysis was performed using deconvolved spike rates.

To exclude redundant neurons due to fluorescence signal bleed-through between two areas or between two neighboring depths, we removed neurons that were highly correlated neighboring neurons, using similar criteria as previously described^51^. We defined potential duplicated neuron pairs as: (1) spike rate correlation above 0.5; (2) lateral distance between centroids below 5 µm regardless of depths; (3) appeared in adjacent imaging depths in the same imaging area (signal bleed-through in the same area from adjacent imaging planes), or appeared in the same imaging depths in different imaging areas (signal bleed-through across areas from the same imaging plane). In these duplicated neuron pairs, we kept the neuron with highest average fluorescence level, and discarded the one with less fluorescence.

Due to the variable length of the choice window, we defined choice window as the 0.5-s time period before the lick that triggered reward window (equivalent to the 0.5-s period before the reward window). We resampled all behavior data to match the calcium imaging rate. Due to slight differences in imaging rate (caused by slightly different pixel numbers), when combining datasets together, we resampled all dataset to a standard 10 Hz rate. These procedures were applied before all the analysis below.

### Responsive neuron analysis

To identify neurons that are responsive to different task phases, we tested the activity level of individual neurons across task windows, for *T*_tone_, *T*_texture_, *T*_choice_, and *T*_reward_. We first denoised the deconvolved traces by a small Gaussian window (3 frames, sigma=1); then, for each neuron *N*_i_ and each task window *T*_j_, we compared its average activity within the window, and generated a baseline distribution by randomly sampling the same number of frames outside of window *T*_j_ and computing average activity, for 100 times. If the activity of neuron *N*_i_ in *T*_j_ was significantly higher than outside of *T*_j_, determined by a one-tailed Wilcoxon Rank Sum test (p<0.05), then we define neuron *N*_i_ as responsive in task window *T*_j_.

To identify neurons that are discriminative for a specific task variable (tone, texture, choice, and reward), we compared the activity of responsive neurons as defined above for the different values of the task variables. For each task variable (for example texture), there are two potential values *s*_1_ and *s*_2_ (for example, texture 1 and texture 2). We compared the average activity of each neuron within the corresponding task window (texture window in this example) between *s*_1_ trials and *s*_2_ trials, using Wilcoxon Rank Sum test. If the neuronal activity was significantly higher (p<0.05) in *s*_1_ trials, then the neuron was defined as a discriminative neuron with preference for *s*_1_. We performed this procedure for all the four task variables.

### Decoder analysis

We trained four types of linear support vector machine (SVM) decoders: tone, texture, choice, and reward decoders, respectively. Similar to the responsive neuron analysis, each type of decoder was trained with two class labels (texture 1 vs. texture 2, for example), using frame-concatenated spike rate data from the corresponding task window (texture window, for example). A separate decoder was trained for each imaging session and each area (S1, PPC-A, PPC-RL), due to different imaging populations. Only matched pairing trials were used for training decoders.

For each imaging session, we randomly divided all the matched pairing trials into 10 subsets. We trained 10 decoders by excluding one subset at one time, therefore each decoder was trained on 90% of the training set. To avoid overfitting, we regularized the SVM coefficients with ridge (L2) penalty. The regularization term was cross-validated in a log space of 10 parameters from 10^−5^ to 10^1^. The projection strength of each trial was calculated using the decoder that was not trained using this trial. The projection strength at time *t* of a trial was defined as the dot product between the decoder coefficient (without the constant term) and the population spike rate vector at *t*. Shuffled controls were generated by randomizing the neuron identities in the dataset and applying the decoders to shuffled data. To evaluate the decoder performance, the ROC (receiver operating characteristic) curve and the AUC (area under curve) were calculated using standard approaches, and discrimination index (DI) was defined as (AUC-0.5)×2. DI=0 represents chance level; DI=1 represents perfect classification performance.

### CCA analysis

To measure the optimal population correlation between cortical areas, we applied a previously reported method, the canonical correlation analysis (CCA)^16,19^. CCA identifies pairs of dimensions from the imaged neuronal populations in the two cortical areas, such that the correlation between the projected activities onto the dimension is maximized. Given the activity of two neuronal populations, a *n*_x_ × *t* matrix **X** from area 1, and a *n*_y_ × *t* matrix **Y**, where t is the number of time points, and *n*_x_ and *n*_y_ are the number of neurons in each area, CCA identifies in total min(*n*_x_,*n*_y_) pairs of dimensions, and the projection correlation of these dimensions decreases from the first to last. Similar to principal component analysis (PCA), CCA finds a set of projection axes for each area; the difference is that PCA aims at maximizing the variance explained by top axes from **X** and **Y**, independently, while CCA aims at maximizing the projection correlation between the activity matrices **X** and **Y**.

CCA requires a sufficient amount of samples to generate stable solutions^35^. In our experiments, we typically imaged 200-300 neurons in each area, and we recorded 200-400 trials per session. A previous study using simulated datasets has shown that ∼50 samples per variable is required to generate a stable solution^35^. To ensure such condition is met, we first performed PCA to reduce the dimensionality (number of variables) per area, keeping the first 30 principal components (PCs). Overall, the top 30 PCs captured a substantial part of the variance in the dataset (variance explained by 30 PCs: S1 55.0±1.2, PPC-A 59.9±1.4, PPC-RL 53.7±1.5 [mean±SEM]). We aimed at generating a separate CCA model at each time point over the trial time; to further increased the sample number, and to avoid outliers as well as to introduce certain temporal smoothness, we took a sliding time window of 0.5 s (5 frames) for training. Using a smaller time window (0.2 s) yielded more noisy but overall similar results (data not shown). For each time window, we randomly divided the data into 10 subsets, and generated 10 models by leaving one subset out each time. The final correlation is computed as the average from the 10 models.

The stimuli in the task result in co-activation of neurons caused by common inputs. As we were interested in the intrinsic interaction between cortical areas, we analyzed the residual activity by subtracting stimulus-triggered averages of PCs, from eight types of stimulus combination (2 tones, 2 textures, 2 choices). Additionally, we observed performance variability within individual sessions, and to ensure that we were analyzing trials during which mice were engaged in the task, we divided each session into sub-sessions of 20 trials, and used only the sub-sessions with performance rate above 75% for analysis. For tone-only and texture-only conditions, we kept the last 50 trials (out of 100-150 trials) when performance had stabilized. This resulted in 40-400 trials for each session.

Since we reduced the dimensionality of each population to its top 30 PCs, the CCA model generated 30 dimensions with descending inter-areal correlation. To determine the number of significant dimensions, we shuffled the trial correspondence between the two areas 100 times and computed the CCA correlations in the same way as above. Significance threshold was defined as mean + 3 S.D. (standard deviation) of the highest shuffled correlation. CCA dimensions with correlations exceeding the significance threshold were regarded as significant dimensions. Overall, we observed 1-2 significant dimensions across datasets and time points, therefore we focused on the first CCA dimension for analysis.

To analyze top-down and bottom-up interactions, we introduced a negative lag of up to 0.5 s with 0.1-s increment to each area, separately. For top-down interaction, we introduced a negative lag to the PPC data (PPC-A or PPC-RL); for bottom-up interaction, we introduced a negative lag to the S1 data. At each lag, we generated a CCA model, consisting of two separate loading matrices corresponding to the two areas. Then, for each trial type (correct, mismatch-choose-texture, and mismatch-choose-tone), we computed the projection correlation of all trials in this trial type. This approach avoided the potential instability of training the CCA model for each trial type, which does not guarantee enough sample numbers. From here, the lagged correlation map was slightly smoothed with a small Gaussian kernel (3 frames, sigma=1) to reduce noise between temporally consecutive models, and top-down and bottom-up interaction strengths were computed as the average CCA correlations across 0.3 s lags. Using different lags (0.1 s, 0.5 s) gave similar results as shown in Fig. 6. The direction and strength of information was quantified as information flow index (IFI), defined as (bottom-up – top-down)/(bottom-up + top-down). IFI is bounded between -1 and 1; values close to -1 represent top-down dominant information, values close to 1 represent bottom-up dominant information, whereas values close to 0 represent simultaneous or no information transfer.

### Pearson correlation analysis

To verify our results of CCA, we also calculated population correlations using Pearson correlation instead of CCA. The imaging data were processed in the same way as for CCA analysis but population correlation was computed as the Pearson correlation between the flattened residual matrices of the two areas. Shuffled correlation was computed from trial-shuffled residual matrices. Unlike CCA, which found positive correlation in all cases, Pearson correlation resulted in negative values in some cases. For the calculation of IFI, we first normalized the lagged correlation map to be between 0 and 1, then calculated IFI for each trial type.

### Statistical analysis

All statistical analysis was done in MATLAB. In general, Wilcoxon signed-rank test was used for paired samples, and Wilcoxon Rank Sum test was used for non-paired samples. All tests were performed with two-sided hypothesis unless otherwise indicated. Error bars represent mean±SEM. Boxplots indicate the median (center line), 25% and 75% quartiles (box limits), and 1.5 × interquartile range (whiskers).

## Code availability

All data processing and analysis code is available from the corresponding author upon reasonable request.

## Data availability

The data that support the findings of this study are available from the corresponding author upon reasonable request.

## Acknowledgements

We thank Philipp Bethge for managing transgenic mouse lines, Fabian Voigt and Hansjörg Kasper for help with optics, and Martin Wieckhorst for the behavior training software. We also thank Christopher Lewis, Jerry Chen and Jordan Hamm for their feedback on the manuscript. This work was supported by a Sinergia grant from the Swiss National Science Foundation (CRSII5_180316; to F.H.) and a UZH Forschungskredit grant (K-41220-07-01; to S.H.). F.H. received funding from the University Research Priority Program (URPP) ‘Adaptive Brain Circuits in Development and Learning’ (AdaBD).

## Author contributions

S.H. and F.H. conceived the study and designed the experiments. S.H. performed the experiments and analyzed the data. S.H. and F.H. wrote the manuscript.

## Competing interests

The authors declare no competing interests.

## Supplemental information

**Supplemental Figure 1.**
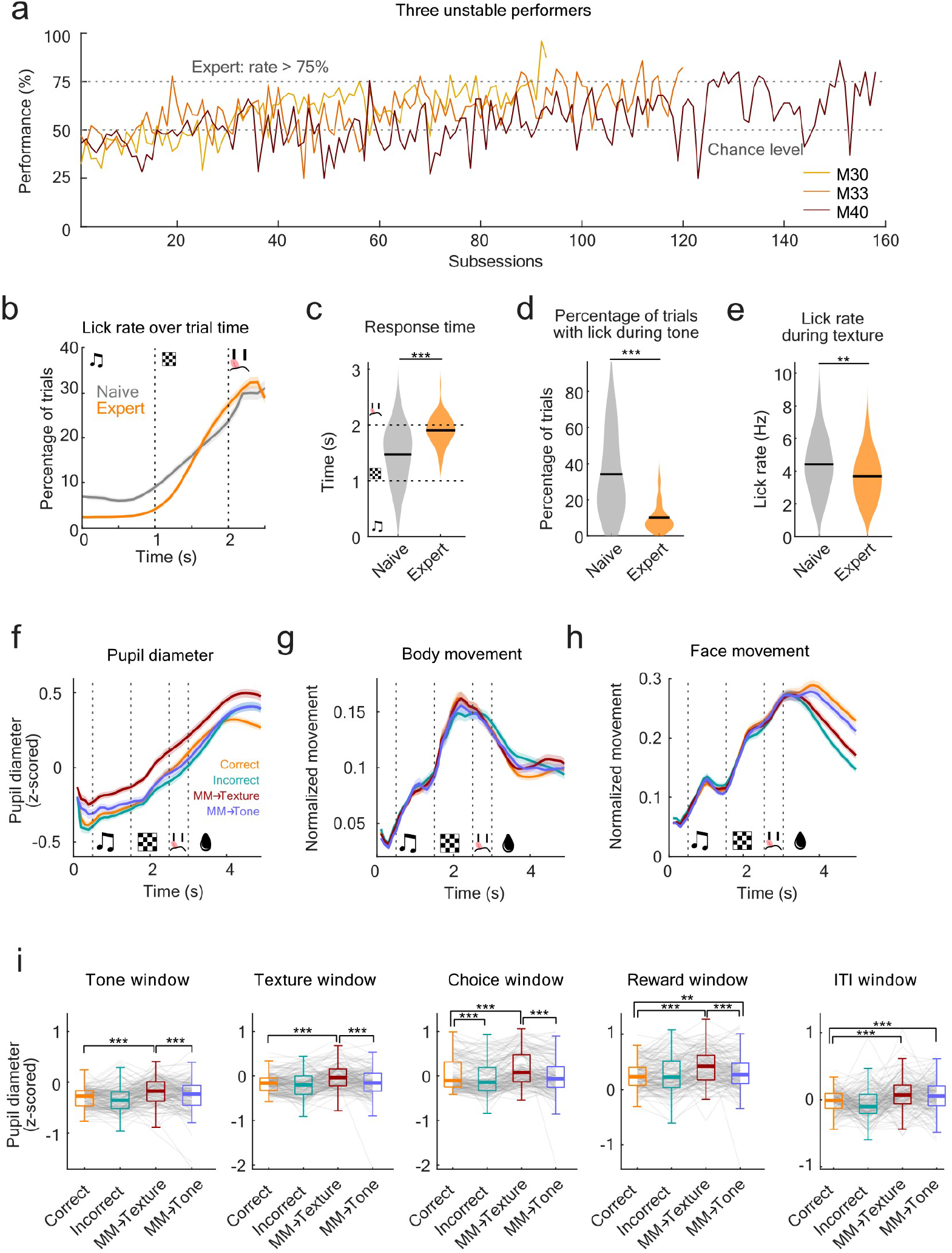
Behavior outliers and movement analysis. (**a**) Each day, all the trials were split into 2-5 subsessions of 100-150 trials. Three unstable performer mice that did not reach expert threshold in daily average all showed expert performance in individual subsessions. (**b**) Lick rate over time for naïve and expert mice. (**c**) Response time of naive (performance<55%, 16 mice, 80 sessions) and expert mice (performance>75%, 13 mice, 74 sessions). (**d**) Percentage of trials with lick during tone on the lick port of final choice for naïve and expert mice. (**e**) Lick rate during texture for naïve and expert mice. (**f**) z-scored pupil diameter in different trial types. (**g**) Body movement (normalized between 0 and 1 within each day) across trial types. (**h**) Face movement (normalized between 0 and 1 within each day) across trial types. (**i**) Statistics of pupil diameter across trial time. (***p<0.001, **p<0.01, *p<0.05, same for all following figures; b-e: Wilcoxon Rank Sum test, i: Wilcoxon signed-rank test; mice and session numbers are the same as Fig. 1).

**Supplemental Figure 2.**
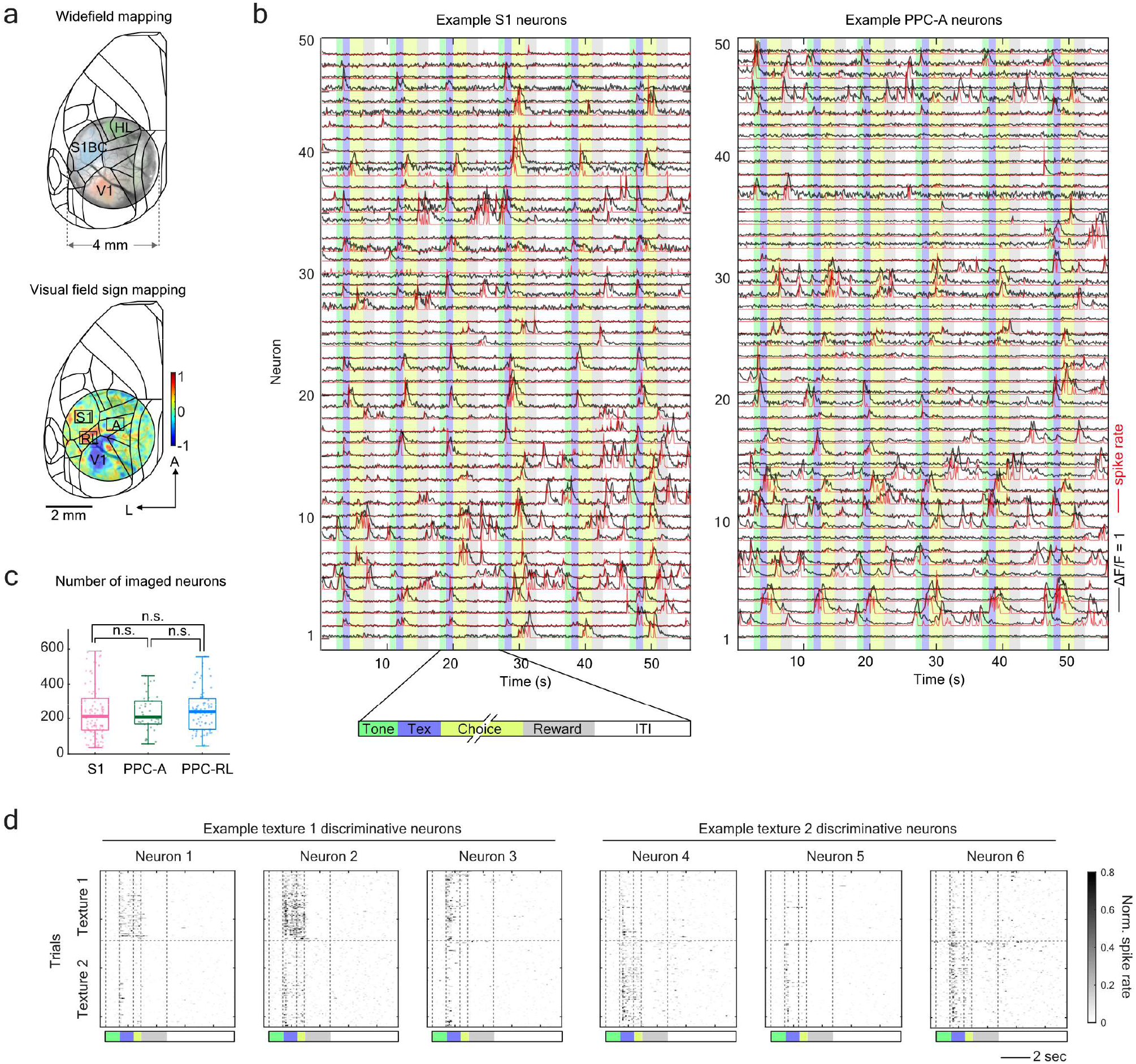
Example traces and additional control experiment statistics. (**a**) The locations of S1, PPC-A and PPC-RL were determined by widefield sensory mapping using whisker, visual and hindlimb stimulation (top) under light anesthesia, as well as visual field sign mapping (bottom). (**b**) Example ΔF/F (black) and deconvolved spike rate (red) of two simultaneously imaged S1 and PPC-A populations. Due to space limitation, only 50 neurons are shown for each area. Colored stripes in the background indicate task windows. Some neurons were silent in the example time period shown in the plot. (**c**) Number of imaged neurons for each area are not significantly different. (**d**) Example single trial activities of texture discriminative neurons. Three example neurons from the same imaging session are shown for each texture preference; choice window was resampled to be the 0.5-s window before the reward window. Trial structure color code is the same as in (b).

**Supplemental Figure 3.**
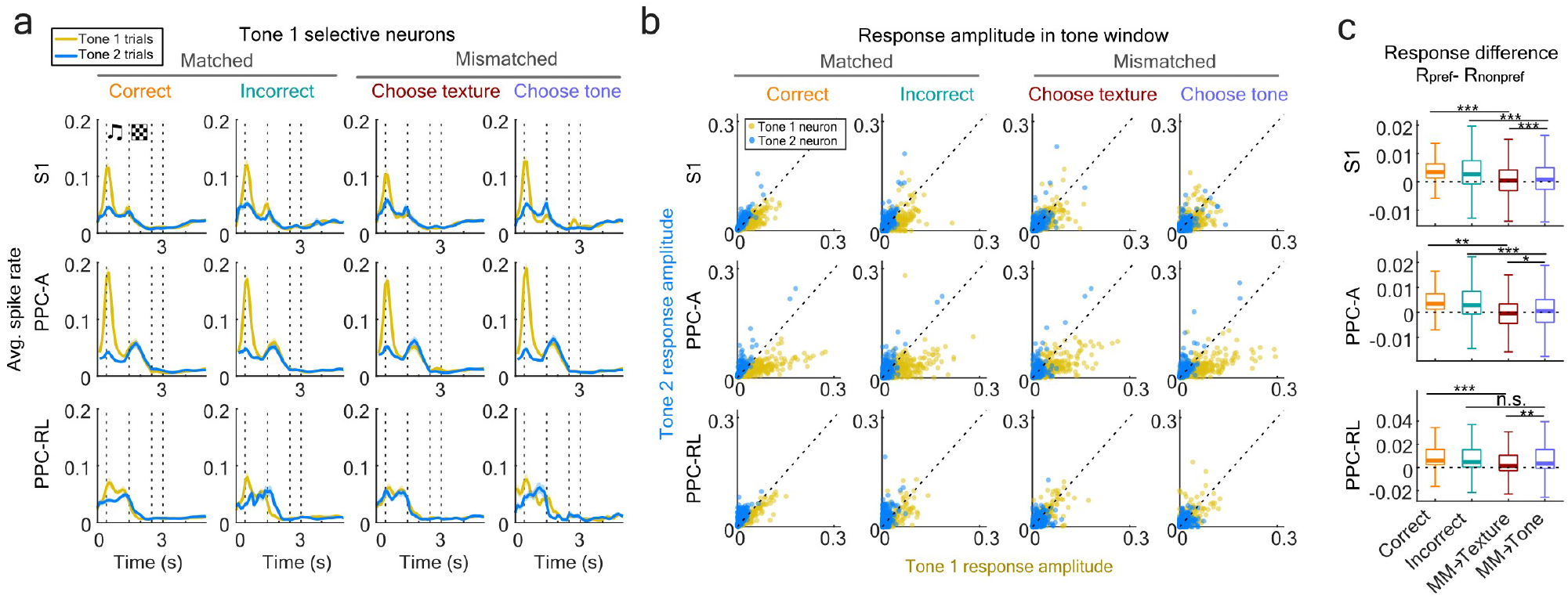
Response amplitude of tone-discriminative neurons. (**a**) Averaged normalized spike rate of tone 1 discriminative neurons in S1, PPC-RL and PPC-A, in matched and mismatched trials. The spike rate of each neuron was normalized to be between 0 and 1 within each session. (**b**) Response amplitude to tone 1 and 2 of all tone discriminative neurons, during tone window, across areas and trial types. (**c**) Selectivity index of texture discriminative neurons during texture window, in different trial types. Significance level was determined from shuffled data where the trial labels were shuffled. (Wilcoxon Rank Sum test; S1: 130 neurons; PPC-A: 179 neurons; PPC-RL: 105 neurons)

**Supplemental Figure 4.**
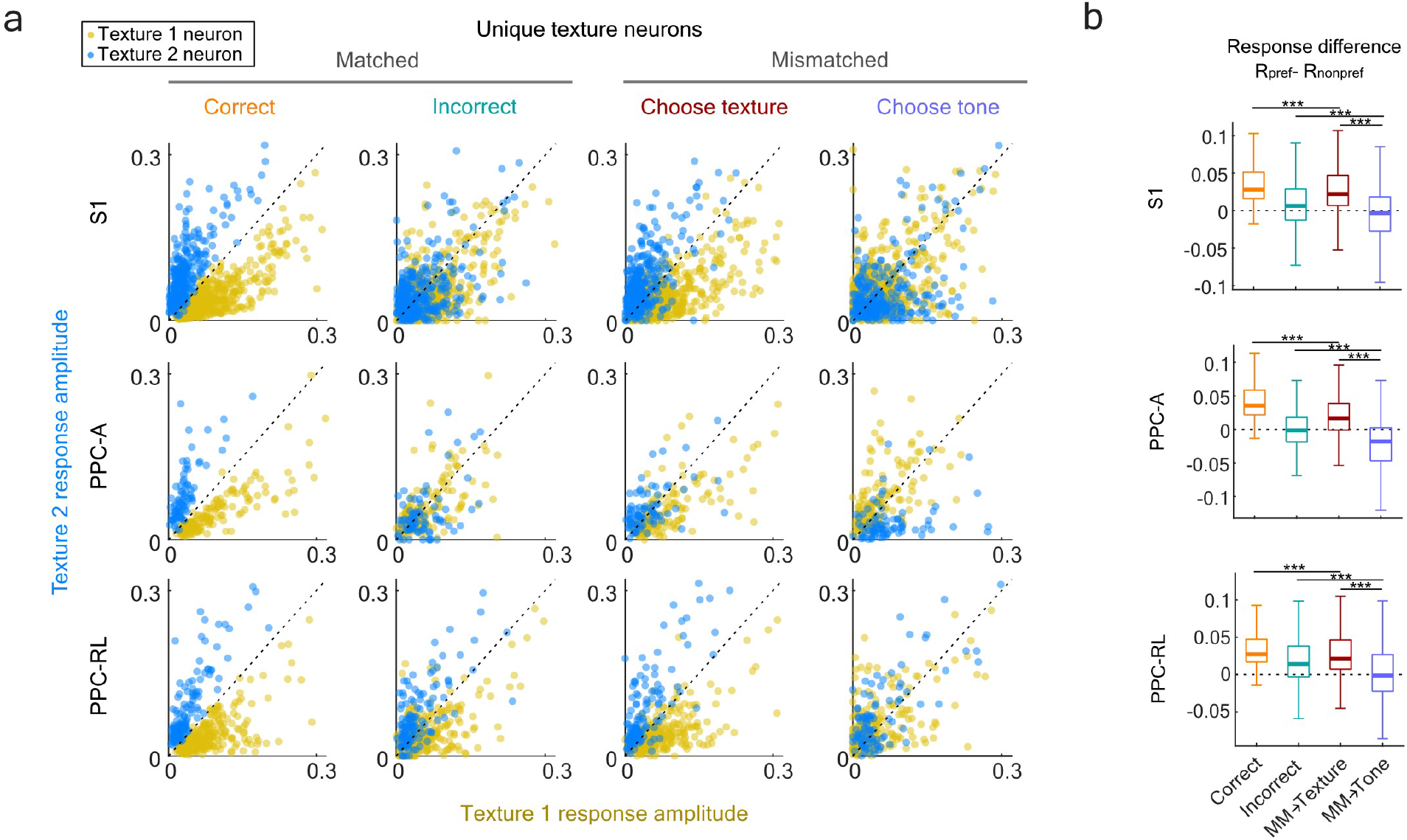
Response amplitude of texture-discriminative neurons that were not choice-responsive. (**a**) Response amplitude to texture 1 and 2 of all texture discriminative neurons that were not choice-responsive, during texture window, across areas and trial types. Each dot represents the response of one neuron in one imaging session. (**b**) Selectivity index of texture discriminative neurons during texture window, in different trial types. Selectivity index was calculated as the difference between the average response to the preferred texture and the average response to the nonpreferred texture. (Wilcoxon Rank Sum test; S1: 1275 neurons; PPC-A: 248 neurons; PPC-RL: 514 neurons)

**Supplemental Figure 5.**
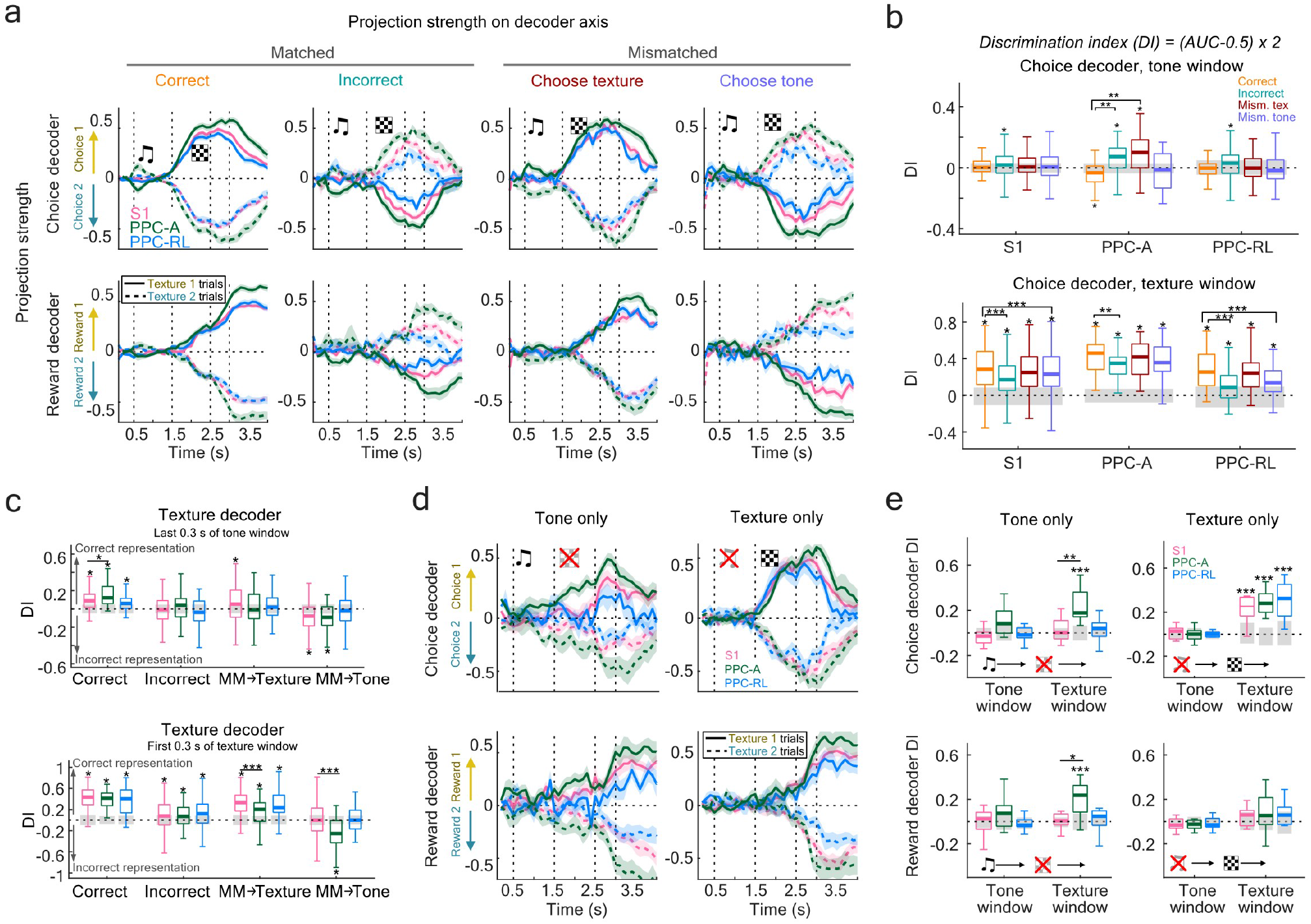
Choice and reward encoding in S1 and PPC. (**a**) Neuronal population encoding of choice (top panels) and reward (bottom panels). Line colors indicate area identity; solid and dash lines indicate texture identity of the trial. (**b**) Discrimination index (DI) of the choice decoder and reward decoder, in the tone window and texture window, separately. Stars above each box indicate the significance with shuffled data (gray bars) where neurons identities were shuffled, while trial and time correspondence were kept the same. Stars across boxes indicate comparison between trial types. (**c**) DI of texture decoder before texture onset (last 0.3 s of tone window, top panel) and after texture onset (first 0.3 s of texture window, bottom panel). (**d**) Neuronal population encoding strength of choice (top panels) and reward (bottom panels) in single modality experiments. (**e**) DI of choice decoder and reward decoder in single modality experiments, in tone window and texture window, separately.

**Supplemental Figure 6.**
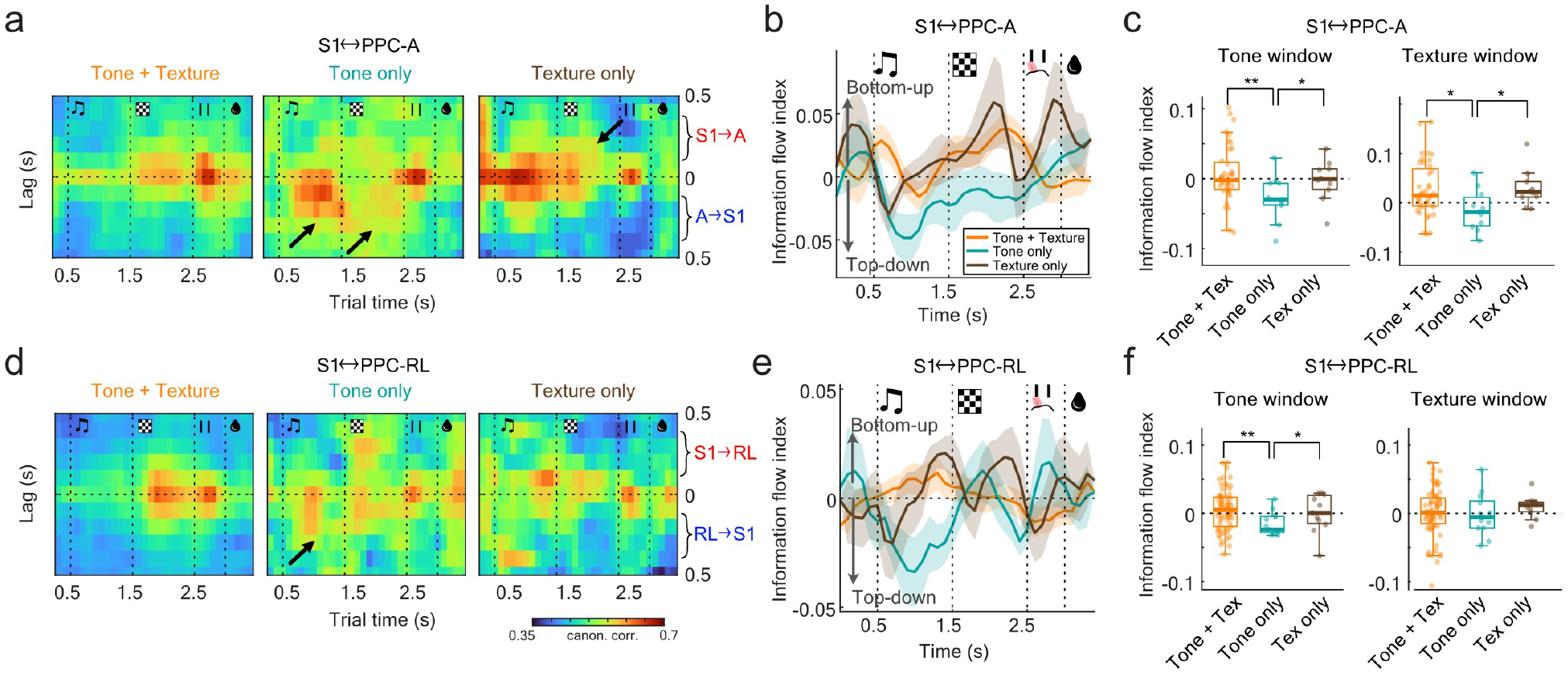
Interaction between S1 and PPC in single modality trials. (**a**) Lagged canonical correlation between S1 and PPC-A averaged across all sessions, for matched stimuli (tone-texture), tone only, and texture only conditions. Note the stronger top-down (A to S1) interaction in tone only condition, and stronger bottom-up (S1 to A) interaction in texture only condition. (**b**) Information flow index quantified from (a). (**c**) Quantification of information flow index between S1 and PPC-A. (**d**) Averaged lagged canonical correlation between S1 and PPC-RL. Note the slightly stronger top-down (RL to S1) interaction during tone window in tone only condition. (**e**) Information flow index quantified from (d). (**f**) Quantification of information flow index between S1 and PPC-RL. (One-sided Wilcoxon Rank Sum test)

**Supplemental Figure 7.**
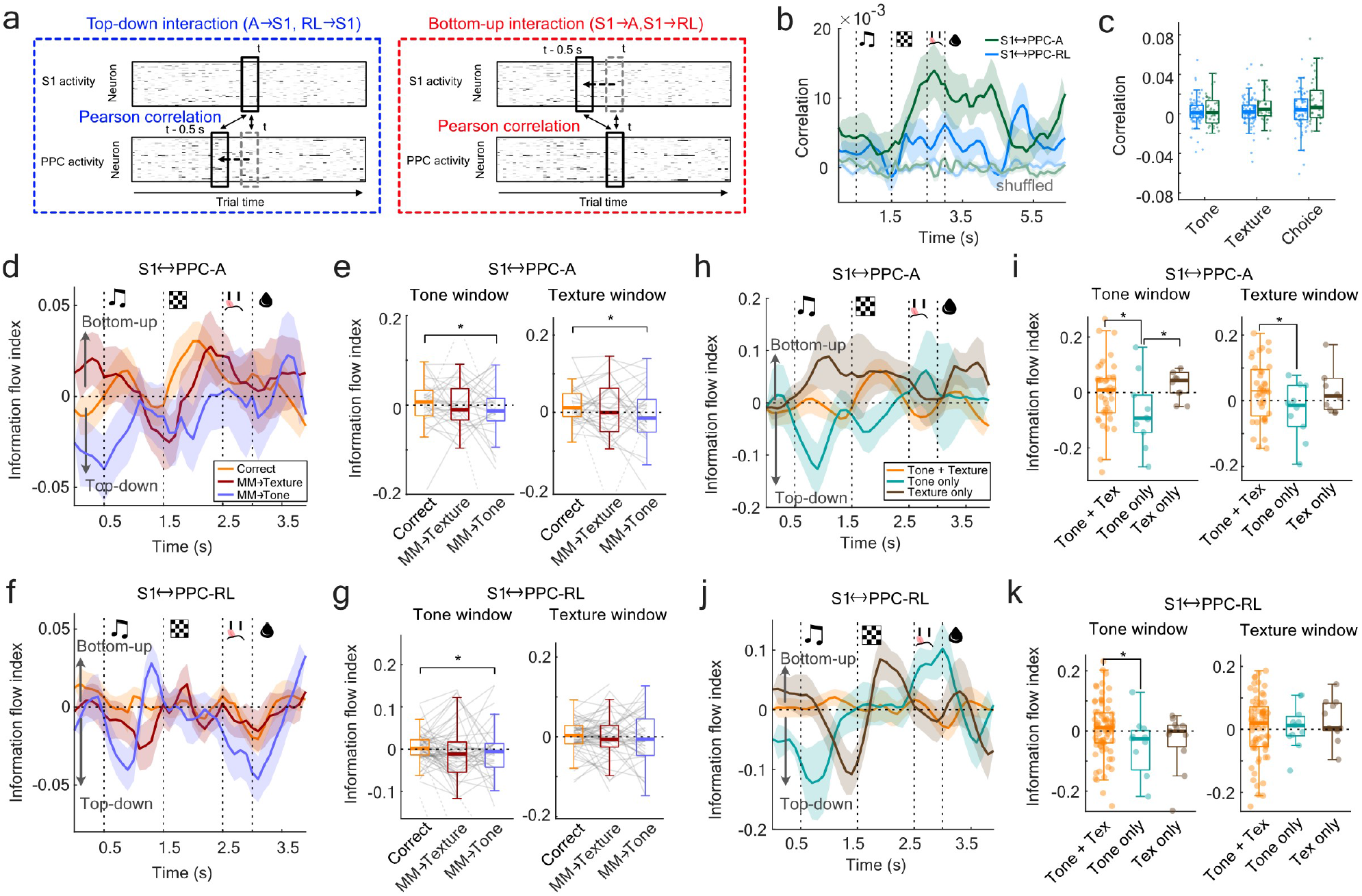
Top-down and bottom-up interaction analysis with Pearson correlation. (**a**) Top-down and bottom-up interaction strength were evaluated in the same way as Fig. 6a, except that the correlation was calculated using Pearson correlation instead of CCA. (**b**) Population correlation between S1 and PPC-A (green), and S1 and PPC-RL (blue). (**c**) S1-PPC_A_ showed slightly stronger interaction than S1-PPC_RL_. (**d**) Information flow index (IFI) of S1-PPC_A_ interaction using Pearson correlation. (**e**) Quantification of (d). (**f**) Information flow index of S1-PPC_RL_ interaction using Pearson correlation. (**g**) Quantification of (f). (**h**) Information flow index of S1-PPC_A_ interaction in single modality conditions. (**i**) Quantification of (h). Consistent with CCA analysis, tone only condition led to stronger top-down information flow during tone and texture windows. (**j**) Information flow index of S1-PPC_RL_ interaction in single modality conditions. (**k**) Quantification of (j). Consistent with CCA analysis, tone only condition led to stronger top-down information flow during tone. (One-sided Wilcoxon signed-rank test)

## Notes

### Competing Interest Statement

The authors have declared no competing interest.

